# MicroRNA-mRNA Gene Regulatory Circuitry Orchestrates the Epigenetic and Transcriptional Reprogramming from Pluripotency to Murine Mammary Epithelial stem-like-Cells lactogenic differentiatio

**DOI:** 10.64898/2025.12.29.696863

**Authors:** Rakhee Nayak, Netrika Tiwari, Sharmistha Chaitali Ghosh, Srinivas Kethavath, Dhammapal Bharne, Sailu Yellaboina, Sreenivasulu Kurukuti

## Abstract

MicroRNAs (miRNAs) are small non-coding RNAs that post-transcriptionally regulate gene expression via mRNA degradation or translational repression. While miRNAs are established regulators of development and differentiation, their role in the dynamic mRNA transitions during lactogenic differentiation of mammary epithelial cells remains incompletely characterized. In this study, we profiled miRNA expression across four distinct developmental states: embryonic stem cells (ESCs), and three progressive stages of HC11 murine mammary epithelial differentiation, including undifferentiated growth factor-maintained cells (HC11-N), glucocorticoid-primed cells (HC11-P), and prolactin-induced lactogenic cells (HC11-PRL). By integrating these signatures with parallel mRNA expression profiles, we identified stage-specific miRNA–mRNA regulatory axes. Our analysis revealed miRNAs coordinate networks of transcription factors and epigenetic regulators to shape the lactogenic transcriptome. While ESCs and undifferentiated HC11 cells were characterized by pathways associated with pluripotency and oncogenic signalling, the pregnancy-like ‘priming’ stage showed enrichment in, neurotrophin signalling. Crucially, the lactation-like state was dominated by PI3K–Akt and mTOR signalling— critical drivers of epithelial proliferation and milk synthesis. Furthermore, the integration of miRNAs with transcriptional and epigenetic regulators highlighted the modulation of prolactin signalling and extracellular matrix–receptor interactions during lactation. This study provides a comprehensive map of the miRNA-associated post-transcriptional landscape of lactogenesis, offering a mechanistic framework for understanding mammary gland development and associated pathologies.

## Introduction

The transformation of a single-cell zygote into a complex multicellular organism depends on precisely coordinated, spatiotemporal gene regulation that drives cell proliferation, lineage commitment, differentiation, and organogenesis (Davidson, 2010; Levine & Tjian 2003). In adult organisms, differentiated cells continuously integrate intrinsic and extrinsic cues to fine-tune gene expression programs, which are orchestrated through multilayered regulatory hierarchies encompassing pre-transcriptional, transcriptional, and post-transcriptional mechanisms. Post-transcriptional epigenetic regulation is particularly vital in non-proliferating differentiated cells, enabling rapid and reversible modulation of gene expression in response to environmental or physiological stimuli (Bartel, 2018; Schwanhäusser et al., 2011). Compared to transcriptional regulation, post-transcriptional control offers the advantage of faster response kinetics, facilitating the immediate fine tuning of the proteome independent of *de novo* transcription.

Among post-transcriptional regulators, microRNAs (miRNAs) constitute a prominent class of small non-coding RNAs that modulate gene expression by directing target mRNA degradation or translational repression during development, lineage specification, and cellular differentiation (Stefanie et al., 2009). The mouse genome contains approximately 2,000 annotated microRNA genes (Cardoso-Moreira et al., 2019), which are evolutionarily conserved and typically 21–26 nucleotides in length. miRNAs predominantly bind to complementary sequences within the 3′ untranslated regions (3′UTRs) of target mRNAs (Confuorti et al., 2024). While perfect complementarity between a miRNA and its target mRNA generally facilitates direct RNA cleavage, the more prevalent mode of action in mammals involves partial complementarity, which inhibits translation or promotes destabilization through deadenylation. Notably, a single miRNA can target hundreds of distinct transcripts, and conversely, individual mRNAs may be co-ordinately regulated by multiple miRNAs in a cell- or tissue-specific manner. Current estimation suggests that over 60% of mammalian protein-coding genes are under selective pressure to maintain miRNAs binding sites (Li et al., 2009). Many miRNAs exhibit stage-specific or tissue-specific expression patterns that drive developmental transitions, with documented regulatory roles in adipogenesis (McGregor et al., 2011), hematopoietic stem cell differentiation (Bissels et al., 2012), and cardiomyocyte lineage specification (Malizia et al., 2011).

Previous transcriptome-wide studies profiling mRNA expression in embryonic stem cells (ESCs) and during the differentiation of HC11 mammary epithelial cell (MEC) have revealed extensive, hormone-dependent, and stage-specific remodelling of global transcriptomic networks (Wang et al., 2009; Williams et al., 2009; Perotti et al., 2009; Sornapudi et al., 2018). These findings underscore the critical role of dynamic gene regulatory programs in driving mammary epithelial specification and functional maturation. However, despite increasing evidence supporting miRNA-mediated post-transcriptional control in developmental processes, the specific miRNA-mediated post-transcriptional mechanisms that orchestrate the divergent transition of pluripotent mouse embryonic stem cells and mammary epithelial lineage remains insufficiently characterized.

The mammalian mammary gland is a highly specialized secretory organ that defines the class Mammalia, distinguishing it from all other vertebrates. Its development is primarily a postnatal process, characterized by coordinated cycles of epithelial proliferation, differentiation, and apoptosis throughout adult life. These dynamic morphological and functional transitions are tightly regulated by systemic hormones and local growth factor signalling networks. In mice, mammary gland morphogenesis initiates at approximately embryonic day 10 (E10) with the formation of a rudimentary epithelial bud (Wu,D. et al., 2022). During puberty, ductal expansion proceeds via the invasion, branching, and proliferation, leading to the formation of terminal end buds (TEBs) that progressively infiltrate the mammary fat pad. Each TEB contains an outer myoepithelial cell layer and an inner luminal epithelial compartment, establishing the structural foundation for subsequent functional differentiation.

During pregnancy, synergistic signalling from prolactin and progesterone stimulate extensive ductal remodelling, including the formation and elongation of alveoli and secondary and tertiary branches (Diana et al., 2022). Following parturition, declining progesterone levels triggers the onset of lactogenesis, driving milk protein synthesis and secretion within the alveolar lumen. Upon weaning, the cessation of suckling cues—coupled with decreased prolactin and increased estrogen signalling—induces massive apoptosis within the alveolar epithelium during involution, ultimately restoring the gland to a quiescent pre-pregnancy state (Confuorti et al., 2024). Consequently, the mammary gland serves as an unparalleled biological model for studying proliferation, lineage commitment, differentiation, and apoptosis outside of embryogenesis, thus enabling dissecting the gene regulatory mechanisms during adult tissue development and remodelling.

The HC11 mammary epithelial cell line, a prolactin-responsive clone derived from the murine COMMA-1D (Balb/c) mouse mammary epithelium, provides a robust in vitro system to dissect these regulatory events (Sornapudi et al., 2018). HC11 cells exhibit bipotent stem-like characteristics and are maintained in the presence of insulin and epidermal growth factor (EGF), representing the undifferentiated, proliferative state (N stage). Upon glucocorticoid-mediated priming (P stage), HC11 cells undergo lineage commitment toward luminal and myoepithelial identities and subsequently acquire terminal alveolar characteristics in response to prolactin (PRL stage), producing milk proteins such as whey acidic protein (Wap) and caseins.

Building upon our previous characterization of steady-state mRNA dynamics across ESCs and HC11 N, P, and PRL states of lactogenic differentiation and demonstrated that thousands of transcripts, including transcription factors and epigenetic regulators, are dynamically regulated during this progression. To investigate post-transcriptional regulation during lactogenic differentiation, we performed comprehensive high-throughput miRNA sequencing across ESCs and HC11 N, P, and PRL stages. Differential expression analysis between ESC–HC11 N, HC11 N–P, and HC11 P–PRL transitions enabled identification of miRNAs and signalling pathways associated with distinct developmental phases. These analyses offer mechanistic insight into the potential roles of miRNAs in orchestrating post-transcriptional gene regulation in murine HC11 mammary epithelial stem-like cells during lactogenic differentiation.

## Results

### Differential expression of microRNAs in murine ESC and mammary epithelial stem-like cells (HC11) undergoing lactogenic differentiation

ESC and HC11 MECs were cultured in presence of EGF (HC11-N), Glucocorticoids (HC11-P) and Prolactin hormone (HC11-PRL) (Fig 1A) in duplicates and were subjected to microRNA isolation and high throughput Illumina sequencing (Sup Fig 1A). MicroRNA data analysis show good correlation between replicates (Sup Fig B & C). Both replicates’ data was mixed and assessed for its abundance (NC) ≥10) in four cellular stages by using MiRDeep2 tool. A total of 388 miRNA genes were found to be expressed in mESC and 122, 142 and 114 miRNAs respectively in HC11-N, P and PRL stages (Fig. 1B). Among them, 249 miRNAs were found to be uniquely expressed in ESC, 2 in HC11(N) and 2 in (P) state (Fig. 1C). Interestingly, 97 microRNAs were found to be expressed among all the cellular stages albeit with variable expression levels (Fig. 1C). To identify differentially expressed miRNAs between ESC-N, N-P and P-PRL stages, MiRDeep2 tool was used to derive differential expressed miRNAs. MiRNAs with Log2 fold change >1, p-value <0.01 were considered as significantly upregulated and Log2 fold change <1 and p-value <0.01 as downregulated. We found an upregulation of 248 miRNAs between ESC-HC11 (N), 51 miRNAs from HC11 (N)-(P). It is interesting to note that only 4 miRNAs were upregulated from HC11-P-PRL stages (Fig. 1B). We also observed down-regulation of 135 miRNAs between ESC-HC11-N and 29, between HC11-N-P stages and found none between HC11-P-PRL stages. To identify key stage-specific miRNAs, the highly expressed that include common and individual miRNAs (Fig. 1D), uniquely expressed (Fig. 2A) and differentially up/down regulated miRNAs were short-listed (Fig. 2B-C). Some of these miRNAs include mmu-miR-122-5p, mmu-let-7a-1-3p, mmu-let-7c-2-3p and mmu-miR-19b-3p were specific for HC11 (N) stage, mmu-miR-155-5p, mmu-miR-1a-3p and mmu-miR-149-5p specific for HC11 (P) stages and mmu-miR-293-5p for HC11 (PRL) stage were validated through real-time PCR (Fig. 2F) and found to be in agreement with respective miRNA-seq datasets (Fig. 2E). Subsequently, we have used these data to dissect complex gene regulatory networks governing the stage specific transitions during lactogenic differentiation of HC11 MECs.

**Fig. 1.**
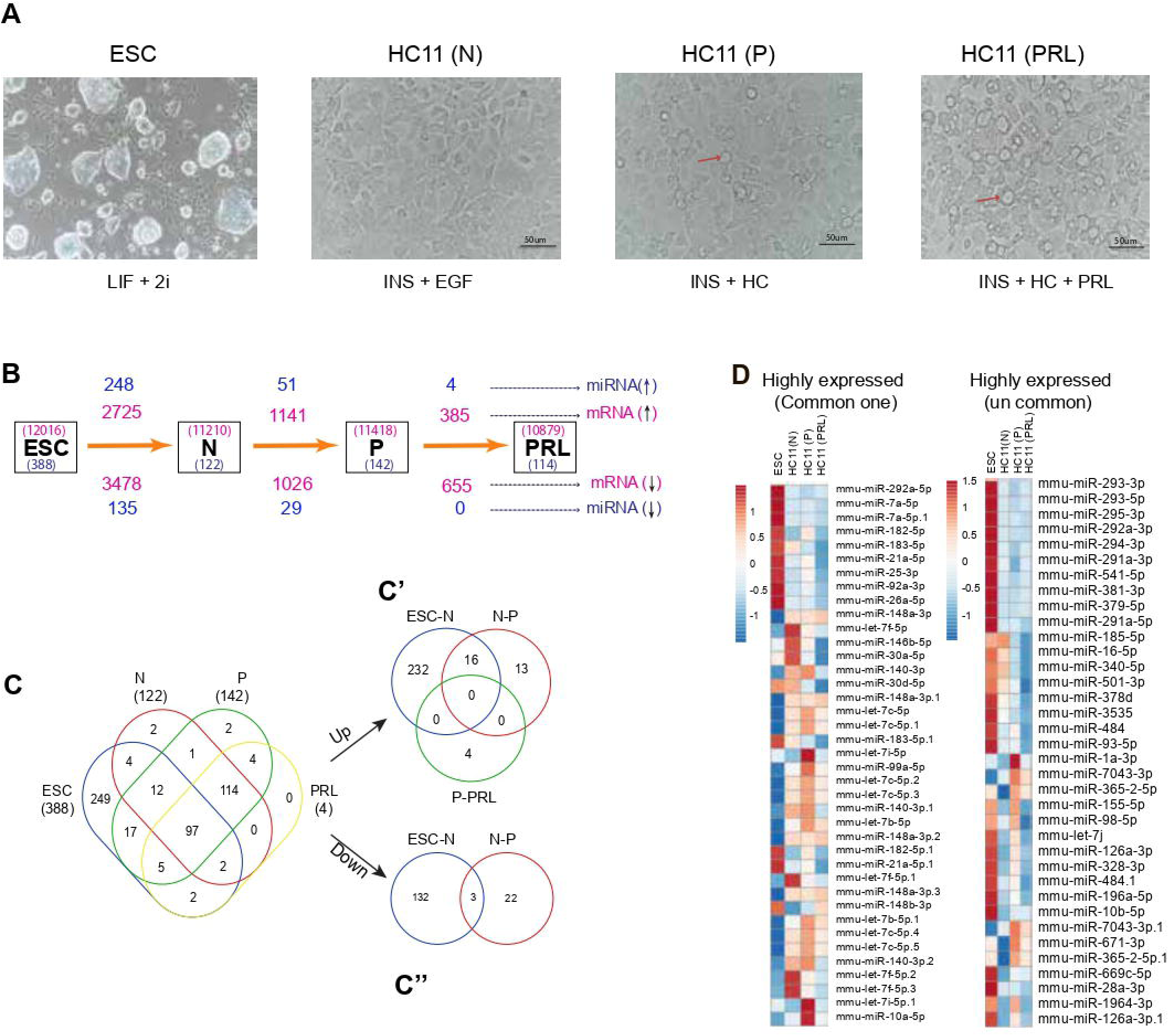
Comprehensive profiling and validation of miRNA-seq. Dataset of ESC and HC11 MECs lactogenic differentiation: (A) Bright field microscopic image of ground state Embryonic stem cell (ESC) cultured in presence of LIF and 2i, undifferentiated HC11 (N) cells at confluent stage in presence of EGF and Insulin, differentiated HC11 (P) cells in presence of Insulin and HC, and differentiated HC11(PRL) cells in presence of Ins, HC, and Prolactin. The red arrow marks represent the formation of mammospheres. Scale bar represents 50μM. (B) Flow chart representing the number of expressed mRNA (in pink) and miRNAs (in blue) in ESC, HC11(N), (P), and (PRL) cell stages, and differentially regulated mRNA and miRNAs in between ESC-HC11(N), (N)-(P), and (P)-(PRL) cell stages. (C) Venn diagram representing unique and overlapping expressed miRNAs (NC≥10) in ESC, HC11(N), (P) and (PRL) cell stages, (C’) differentially upregulated (Log2 Fold change ≥ 1), (C’’) downregulated (Log2 Fold change ≤ −1) miRNAs between ESC-HC11(N), (N)-(P) and (P)-(PRL) cellular states. (D) Heatmap representing expression status of stage-specific miRNAs in HC11(N), (P), and (PRL) cell stages. (E) Bar graph representing normalized counts of miRNA-seq data of stage-specific miRNAs. (F) Real-time PCR validation of miRNAs for HC11 (N), (P), and (PRL) cell stages. Data were analyzed using the −2^ΔΔ^CT method. *Rnu6* and HC11(N) were considered as reference and sample control, respectively. Bar graphs were generated using GraphPad Prism software with a one-way ANOVA significance test. MECs: Mammary epithelial cells; LIF: Leukemia inhibitory factor; 2i: two inhibitors; Ins: Insulin; HC: hydrocortisone, N: Normal, P: Primed, PRL: Prolactin

**Fig. 2.**
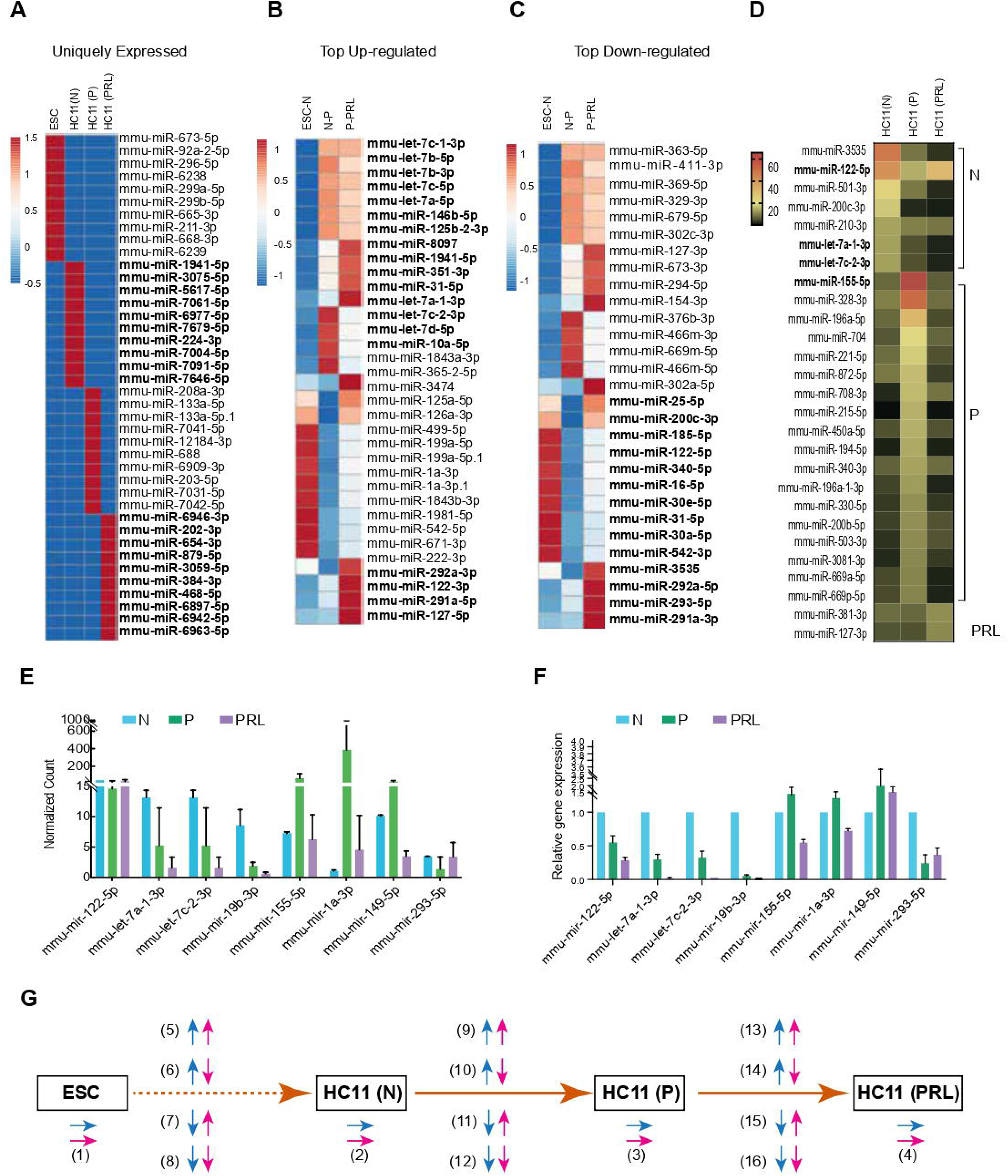
Comparative analysis of uniquely and differentially expressed miRNAs in ESC and HC11 MECs cells undergoing lactogenic differentiation: Heatmap representing highly expressed (Highest value of NC), miRNAs in two replicates of ESC (1&2), N (1&2), P (1&2) and PRL (1&2) (A), exclusively expressed (unique to each stage) (B), Top upregulated (Highest value among Log2 Fold change ≥ 1) (C), and Top downregulated (Lowest value among Log2 Fold change ≤ −1) (D). NC: Normalized count

### MiRNA-mRNA interactome analysis in ESCs

MicroRNAs target expressed mRNAs in ESCs, potentially preventing them from undergoing translation. Using the miRNet tool (Chang et al., 2020), which considers experimentally validated miRNA-mRNA interaction targets, we predicted mRNA pathways that were controlled by miRNAs. In the core of the interactome maps, miRNAs to target more than 25 mRNAs such as mmu-miR-340-5p, 329-3p, 9-5p, 7b-5p, 124-3p, 144-5p, 26a-5p, 466d-5p, 17-5p, 425-5p, 301b-3p, etc. were noticed and were found to be controlling the mRNA of the genes involved in Pathways in Cancer, signaling pathways regulating pluripotency of stem cells, MAPK signaling pathway, Hippo signaling pathway, Axon guidance, Focal adhesion, ErbB signaling pathway, etc. were found to be controlled potentially may not be allowing them to undergo translation (Fig. 3A). Among them, the miRNA:mRNA network of Signaling pathways regulating pluripotency of stem cells (Fig. 3A’) was highlighted.

**Fig. 3.**
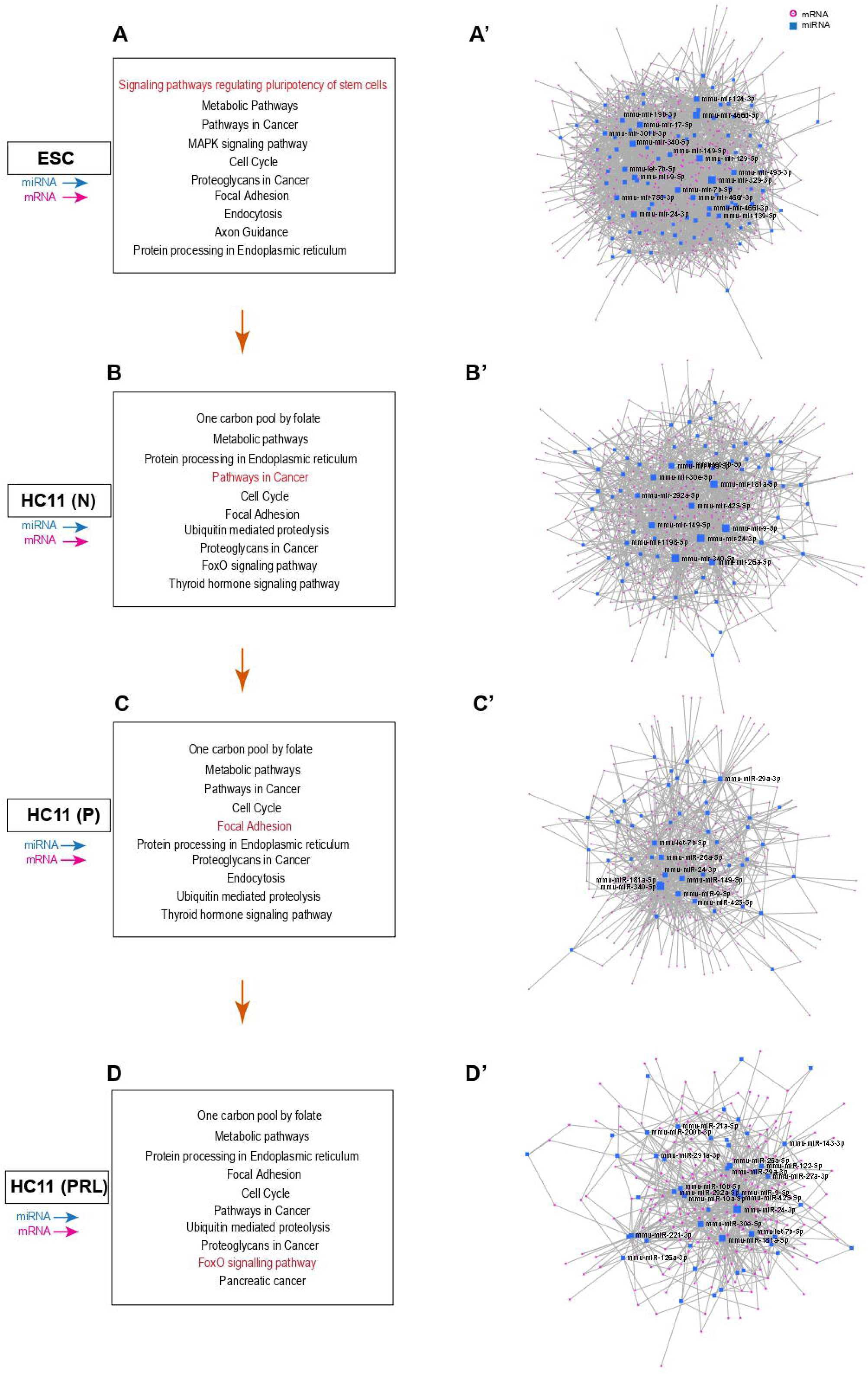
Post-transcriptional control of key gene mRNAs by miRNAs involved in specific pathways in ESC (A), HC11-N (B), P (C), and PRL (D) cell stages by considering expressed miRNAs (NC≥10) and their targets filtered with respective stage-specific mRNAs (FPKM≥ 1). From each cell stage, one of the pathways (red colour) specific miRNA-mRNA interactome networks was highlighted. Signalling pathways regulating pluripotency in stem cells in ESC (A’), pathways in cancer in HC11-N (B’), focal adhesion in HC11-P (C’), and the FoxO signalling pathway in HC11(PRL) cell stages.

### miRNA-mRNA interactome analysis in HC11 (N) MECs

MicroRNAs, expressed in HC11(N) may target expressed mRNAs in HC11 (N) MECs and potentially not allow them to undergo translation. Dissecting the regulatory principles of gene expression in this state is crucial, as dysregulation at this point can lead to the failure of lactogenesis or might skew developmental programs towards carcinogenesis (Wang et al., 2019). MiRNA-mRNA interactome maps were generated considering expressed mRNAs and miRNAs in the MEC (N) state and found that miRNAs such as mmu-mir-340-5p, 9-5p, 181a-5p, 26a-5p, 149-5p, 30e-5p, 425-5p, 24-3p, let-7b-5p, 10a-5p, 292a-5p, 1198-5p, etc regulating mRNAs involved in Pathways in Cancer, Neurotrophins signaling pathway, FoxO signaling Pathway, Focal Adhesion, Apoptosis, HIF-1 signaling pathway, etc. (Fig. 3B&B’). These pathways are known to play an important role in cellular homeostasis, metabolic activity, growth and development and were silenced at HC11 (N) stage of lactogenesis. Here, an enlarged view of the miRNA:mRNA network of Pathways in cancer (Fig. 3B’) is provided. Pathways in Cancer are significantly controlled in all the four stages by many miRNAs, creating a complex miRNA:mRNA network however, interestingly all the networks differ at each stage and composed of distinct miRNAs and mRNAs.

### miRNA-mRNA interactome analysis in HC11 (P) MECs

Expressed microRNAs in HC11(P) target expressed mRNAs in HC11 (P) MECs, potentially may be controlling the rate of translation. The miRNA-mRNA interactome analysis for MEC(P) cells revealed major miRNAs’ hub towards MEC(P) transition were found to be centered around microRNAs such as mmu-mir-10b-5p, 292a-5p, 1198-5p, 122-5p, 155-5p, 1981-5p, 185-5p, 34a-5p, etc. controlling genes involved in pathways such as, Neurotrophins signaling pathway, Chronic myeloid leukemia, HIF-1 signaling pathway, Focal Adhesion, Foxo signaling pathway, MAPK signaling pathway, Axon Guidance, Hippo signaling pathway, ErbB signaling pathway, etc. (Fig. 3C). The miRNA:mRNA network of Focal Adhesion (Fig. 3C’) is highlighted here. Focal Adhesion has an integral role in cell migration and ductal development during lactogenesis. These kinds of pathways may not be getting suppressed instead may be driven under control.

### miRNA-mRNA interactome analysis in HC11 (PRL) MECs

MicroRNAs targeting expressed mRNAs in HC11 (PRL) MECs, potentially may control the translation. Major miRNAs hubs that were identified in MEC(PRL) stages were found to be derived from mmu-mir-9-5p, 181a-5p, 26a-5p, 425-5p, 30e-5p, 24-3p, 7b-5p, 10a-5p, 10b-5p, 292a-5p, 30a-5p, 30c-5p, 451a, etc. controlling mRNAs involved in pathways such as mTOR signaling pathway, PI3K-Akt signaling pathway, Endocytosis, TNF signaling pathway, MicroRNAs in cancer, Wnt signaling pathway, Oxytocin signaling pathway, AMPK signaling pathway, etc. (Fig 3D). Among the above, the miRNA:mRNA network of FoxO signalling pathway (Fig 3D’) was extracted from the main network. The FoxO signaling pathway is vital in mammary gland development, which is essential for the survival and proliferation of mammary cells. This pathway has been significantly controlled in the Prolactin state.

### Differentially Expressed miRNA-mRNA interactome analysis between ESC-HC11 (N) stages

MiRNA regulates genes by mRNA degradation or translational arrest. Keeping these in view, to draw a significant interactome map, differentially upregulated miRNAs were analysed together with differentially downregulated mRNAs and differentially downregulated miRNAs along with differentially upregulated mRNAs. Upregulated miRNAs targeting mRNAs lead to downregulation of their levels, potentially preventing them from being translated in the subsequent stage. To study the downregulated interaction map of HC11(N)_ESC, the upregulated miRNAs and downregulated mRNAs between HC11(N)_ESC were considered, where the list of downregulated mRNAs was filtered out from the total targeted mRNAs. The central hub of this interactome consists of mmu-mir-9-5p, 181a-5p, 340-5p, 149-5p, 7b-5p, 24-3p, 26a-5p, 30e-5p, 10a-5p and 362-5p with more than 50 targets. These miRNAs are mainly associated with pathways like Pathways in cancer, Wnt Signalling Pathway, P53 signalling pathway, signaling pathway regulating pluripotency, etc. (Fig. 4A). Among all, the miRNA:mRNA interaction map of cAMP signalling pathway was shown in Fig. 4A’. This pathway is essential for maintaining ESC self-renewal and has been found to be downregulated during differentiation (Faherty et al. 2007).

**Fig. 4.**
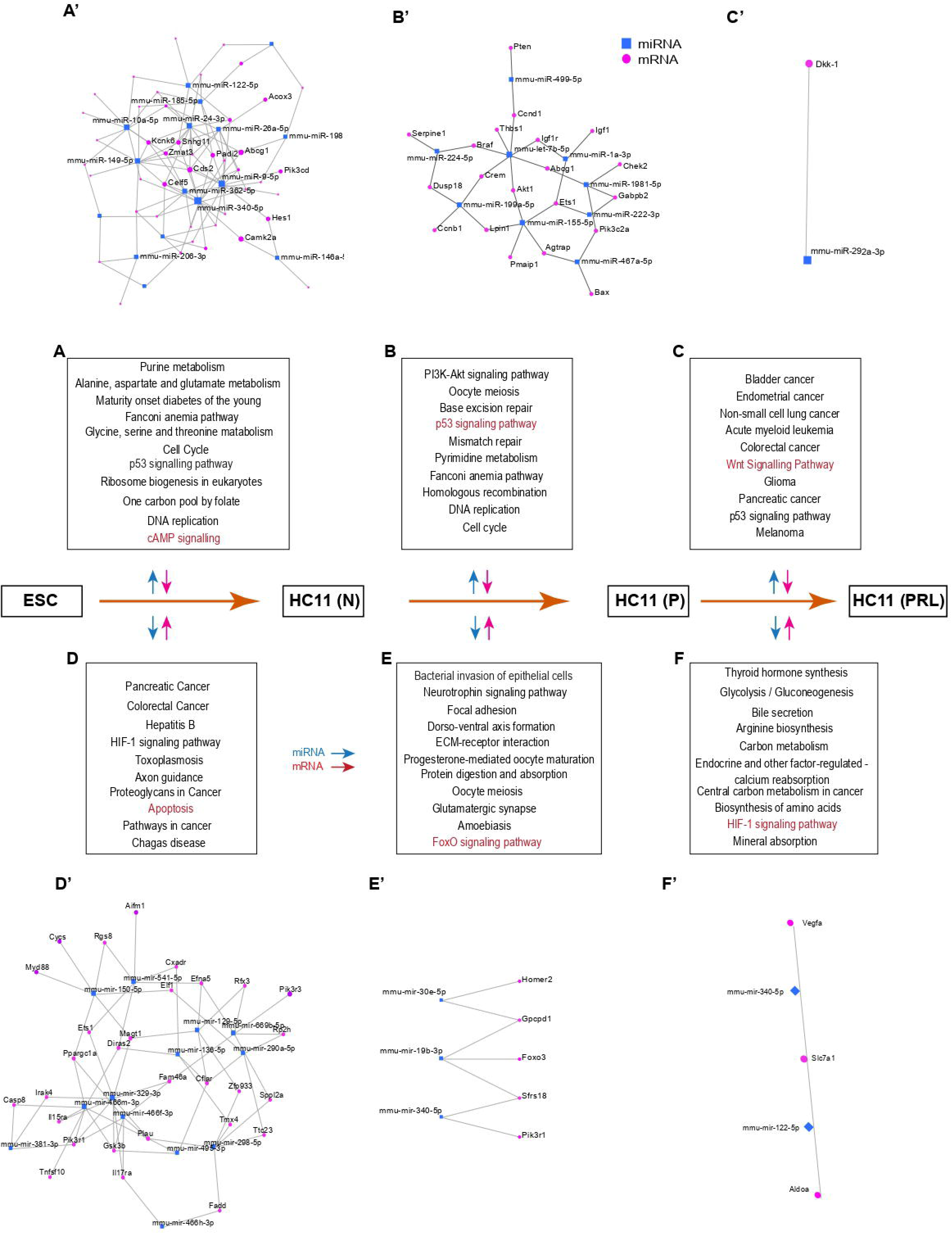
Post-transcriptional control of down or up-regulated mRNAs of genes and their specific pathways by down or up-regulated miRNAs, respectively, during cell transitions from ESC-HC11(N) (A&D), N-P (B&E), and P-PRL (C&F) cellular stages. A representative miRNA-mRNA interactome maps (highlighted in red colour) for Apoptosis (A’), FoxO signalling (B’), HIF-1 signalling pathway (C’), cAMP signalling pathways (D’), P53 signalling pathways (E’), and Wnt signalling pathway (F’) are depicted for each category.

An interaction map was generated by considering upregulated mRNAs and downregulated miRNAs of ESC-HC11(N). Here, we identified several miRNAs that serve as central hubs in the interactome with more than 50 targets like, Mmu-mir-7b-5p, 329-3p, 129-5p, 541-5p, 466f-3p, 136-5p and 466m-3p. This interaction also regulates many key important pathways. Among them, the interaction map of pathways (Hippo signaling pathway, PI3k-Akt Signaling pathway, signaling pathway regulating pluripotency and Focal Adhesion) are highlighted (Fig. 4D) with an interaction map of Apoptosis (Fig. 4D’).

### Differentially expressed miRNA-mRNA interactome analysis between HC11 (N)-(P) stages

Upregulated miRNAs targeting mRNAs lead to downregulation of their levels potentially not allowing them to translate in subsequent stages. The interaction map of upregulated miRNAs and downregulated mRNAs of HC11(P)_HC11(N) did not come up with any significant pathways. However, many miRNAs like mmu-mir-let-7b-5p, 155-5p, 126a-3p, 204-5p, 199a-5p, 328-3p, 1a-3p, 224-5p, 429-3p, 1981-5p, 125a-5p, 467a-5p, 151-3p and 6669c-5p were involved in the downregulation of gene transcripts (Fig. 4B-B’).

To understand the significance of downregulated miRNAs in HC11 (P), we analysed the miRNA-mRNA interaction map of downregulated miRNAs and upregulated mRNAs from HC11(P)_HC11(N) stages. This analysis showed that the primary miRNAs’ hubs cantered around multiple miRNAs such as mmu-mir-340-5p, 19b-3p, 30e-5p, 292a-5p, 185-5p, 203-3p, let-7e-5p and let-7c-5p. This interaction map was involved in controlling many important pathways in primed conditions like Bacterial invasion of epithelial cells, progesterone-mediated oocyte maturation, ECM-receptor interaction and Amoebiasis (Fig. 4E). A significantly upregulated pathway during lactogenic differentiation is FoxO signaling pathway (Fig. 4E’).

### Differentially expressed miRNA-mRNA interactome analysis between HC11 (P)-(PRL) stages

Upregulated miRNAs targeting mRNAs lead to the downregulation of their expression levels. Considering the downregulation map of HC11(PRL)_HC11(P), it did not provide any significant interactions in pathways. The pathways list is obtained by considering only upregulated miRNAs of HC11(Prl)_vs HC11(P) (Fig. 4C). Among the upregulated miRNAs, only mmu-mir-292a-3p and 291a-5p showed interaction with downregulating genes *Dkk-1* and *Ttc14,* respectively (Fig. 4C’).

Likewise, to extract the miRNA-mRNA interaction map of HC11(PRL), we considered upregulated mRNAs between HC11(PRL)_HC11(P) and downregulated miRNAs between HC11(P)_HC11(N). There were no significant differentially downregulated miRNAs between HC11(PRL)_HC11(P), which is why downregulated miRNAs from HC11(P)_HC11(N) were considered, as the transcript levels of miRNAs were maintained throughout the primed and Prolactin states. This interaction map comprises miRNAs like, mmu-mir-340-5p, 19b-3p, 30e-5p, 122-5p and let-7e-5p, which regulate pathways like Cell Cycle, mTOR Signaling pathway, HIF-1 signaling pathway and PI3k-Akt signaling pathway (Fig. 4F). The highlighted HIF-1 signalling network is reported to be involved in lactogenic differentiation during lactation period (Paatero et al., 2014).

### Up-regulated miRNA-mRNA genes interactome analysis during cell state transitions

Many upregulated miRNAs can also target some of the upregulated mRNAs because they are expressed at the same time and share complementarity. The purpose of these interactions may not be to abolish the translation process altogether, but rather to control the expression of genes to some extent in a process aimed at achieving successful differentiation. However, spaciotemporarily in a cellular biochemical system, whether probable interactions exist or not, needs to be validated experimentally. Here, we attempted to identify potential interactions between upregulated miRNA:mRNA through bioinformatic analysis. The miRNA:mRNA network between HC11(N)_ESC showed pathways like Bacterial invasion of epithelial cells, Pathways in cancer, Signaling pathways regulating pluripotency, etc. were targeted (Fig. 5A). Among them, the interaction map of the Hippo signaling pathway is highlighted (Fig. 5A’). The interaction map between HC11(P) and HC11(N) provided pathways like Acute myeloid Leukemia, Adherens Junction, Melanogenesis, etc. (Fig. 5 B-B’). Likewise, the interaction map of HC11(P)_ HC11(Prl) generated only one significant pathway that is the Wnt signaling pathway, targeted by miR-292a-3p (Fig. 5 C-C’).

**Fig. 5.**
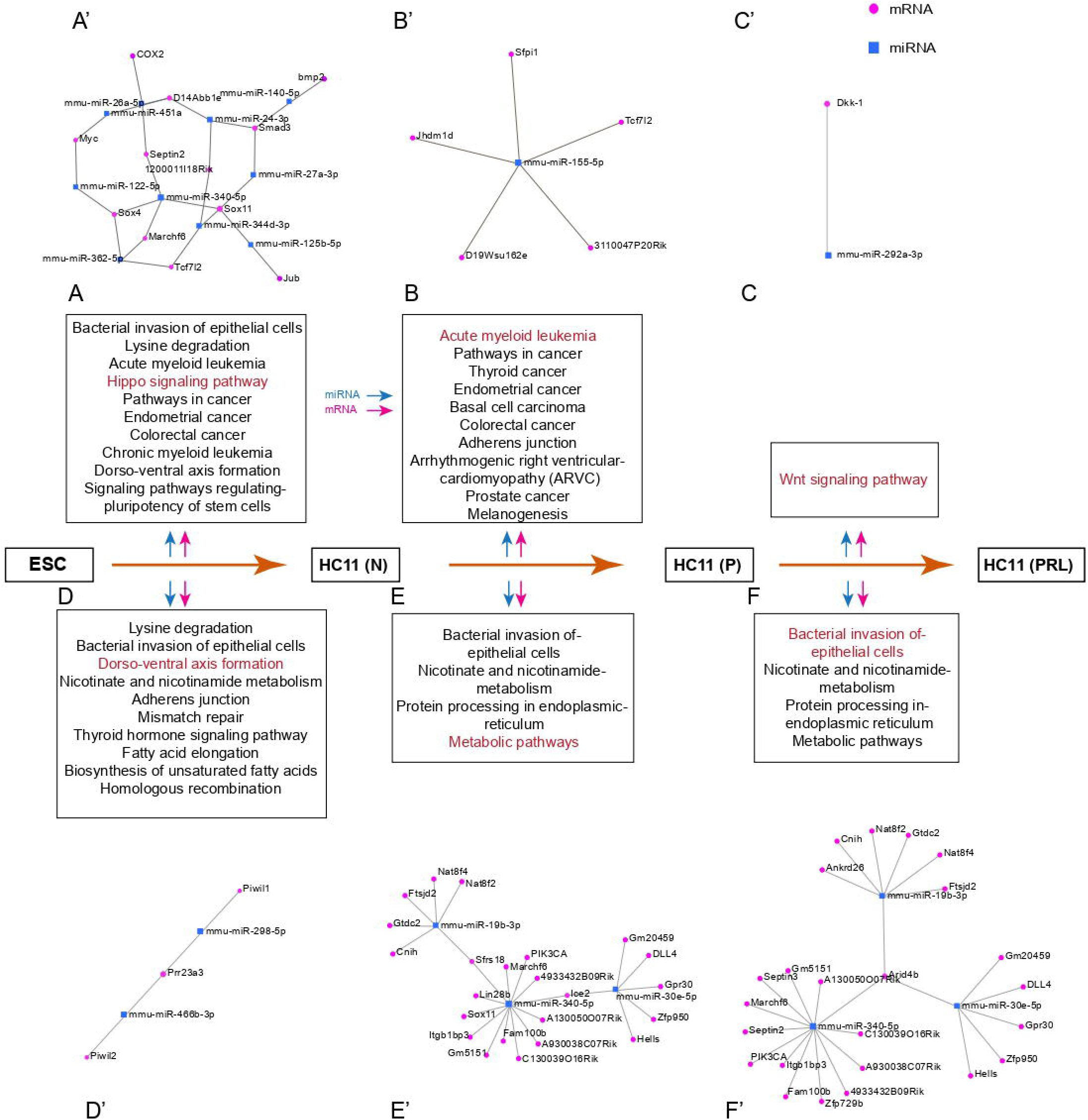
Post-transcriptional control of up and down regulation of mRNA and miRNAs and its specific pathways during cell transitions from ESC-HC11 (N) (A&D), N-P (B&E) and P-PRL (C&F) and its representative miRNA-mRNA interactome. (highlighted in red colour) of Hippo signalling pathway (A’), Acute myeloid leukemia (B’), Wnt signalling pathway (C’), Dorso-ventral axis formation pathway (D), Metabolic pathway (E’) and Bacterial invasion of epithelial cells pathway (F’) are depicted for each category.

### Down-regulated miRNA-mRNA genes interactome analysis during cell state transitions

Similarly, by considering downregulated miRNAs with downregulated mRNAs, potential interaction maps were generated. The miRNA:mRNA network from HC11(N) vs ESC came up with pathways like Bacterial invasion of epithelial cells, Adherens junction, Thyroid hormone signaling, etc. were targeted (Fig. 5D). Among them, the interaction map of the Dorso-ventral axis formation pathway was highlighted (Fig. 5D’). The interaction map of HC11(P)_HC11(N) was linked with pathways like Bacterial invasion of epithelial cells, Metabolic pathways, etc. (Fig. 5 E-E’). In a similar way, the miRNA:mRNA interaction map of HC11(P)_ HC11(Prl) also controlled pathways like Bacterial invasion of epithelial cells, etc. (Fig. 5 F-F’). The same pathway is controlled by different miRNAs at various stages of differentiation, targeting either similar or distinct genes within that pathway.

### miRNA-TF mRNA interactome analysis

Cell state transitions during cell specification and differentiation are governed by transcription factors (TF) and epigenetic regulators (ERs). The potential targeting of TF/ER mRNAs by miRNAs during cell state transition has been investigated. The list of transcription factors (TF) was filtered out from the transcribed gene list of expressed and differentially regulated to proceed with the integration analysis with miRNA. Firstly, we have identified the TFs that are regulated by the miRNAs in a stage-specific manner (Fig. 6A). In ESC, 267 miRNAs were targeted in a total of 1054 expressed TFs. Further, we analysed the targeted pathways and found the pathways like Wnt signaling pathway, Insulin resistance, Hippo signaling pathway, Lysin degradation, MAPK signaling pathway, etc (Fig. 6B’-B). There were 93 miRNAs in the case of HC11(N) that targeted 622 TFs and the pathways being targeted are TNF signaling pathway, Thyroid hormone signaling pathway, Tuberculosis, Adherens junction, Insulin resistance, etc. (Fig. 6C’-C). Similarly, 104 and 85 miRNAs in HC11(P) and HC11(PRL) were targeted in a total of 648 and 540 TFs respectively (Fig. 6D’ and E’). In the case of HC11(P), the pathways being targeted through TFs are Wnt signaling pathway, Estrogen signaling pathway, MAPK signaling pathway, Thyroid hormone signaling pathway, Adherens junction, Prolactin signaling pathway, etc. (Fig. 6D) and in the case of HC11(PRL), the pathways are TNF signaling pathway, Circadian rhythm, Hippo signaling pathway, Notch signaling pathway, Adherens junction, Prolactin signaling pathway, etc. (Fig. 6E). Further, we analysed differentially expressed TFs and targeted miRNAs. The same logic has been applied here, for the list of upregulated TFs between states’ targeted interactions with the downregulated miRNAs and vice versa were visualized. Sixty four downregulated miRNAs that are targeting 133 upregulated TFs and 89 upregulated miRNAs targeting 219 downregulated TFs. During HC11(N) to HC11(P) differentiation, six upregulated miRNAs were identified as targeting 19 downregulated TFs and 12 downregulated miRNAs targeting 33 upregulated TFs. Similarly, in HC11(PRL) vs HC11(P) only one upregulated miRNA has been identified that targets one downregulated mRNA (Fig. 6A).

**Fig. 6.**
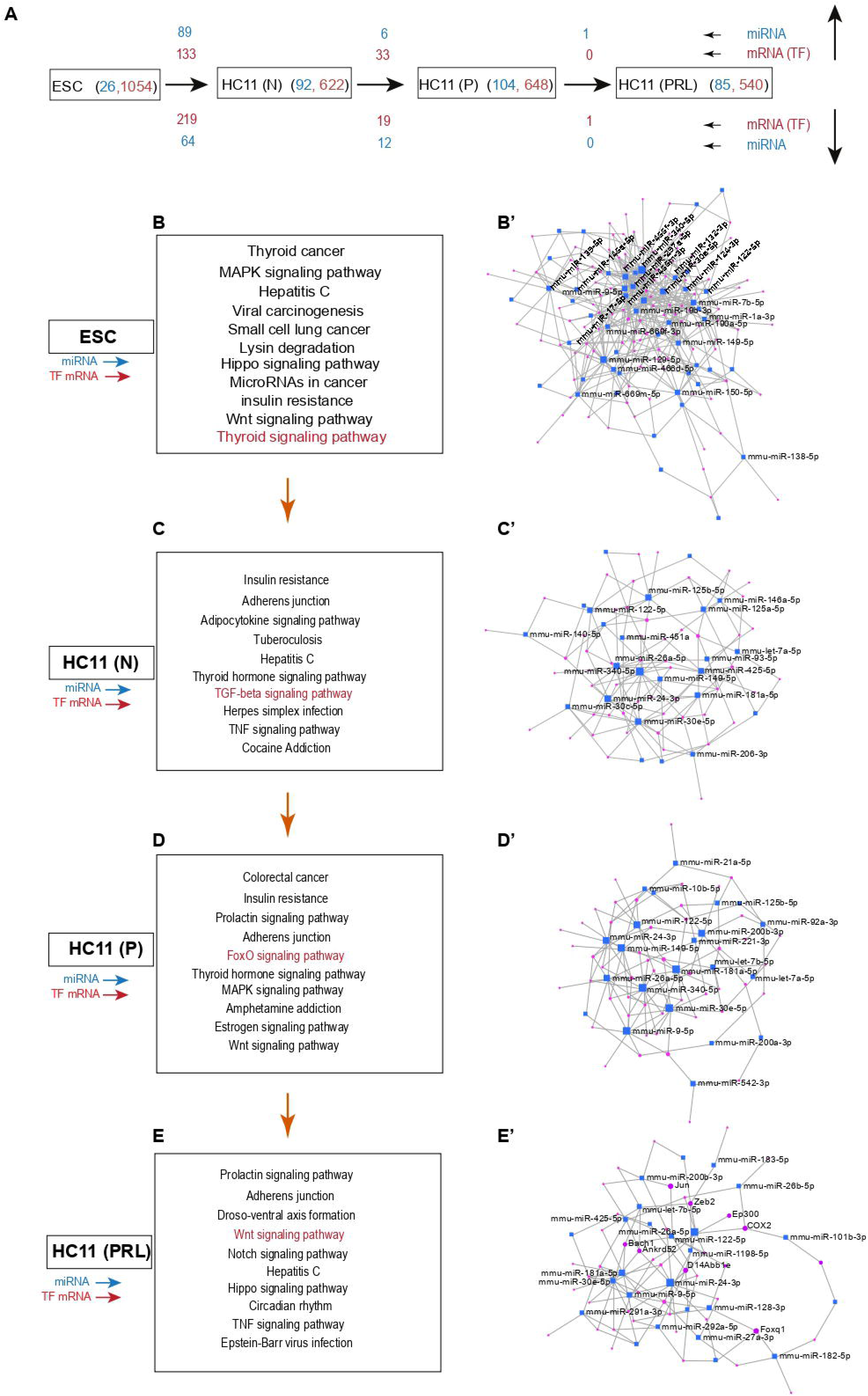
Post-transcriptional control of stage-specific Transcription factor (TF) mRNAs and their specific pathways by miRNAs in ESC, HC11 (N), (P), and (PRL) cell stages. (A) Flow chart representing the total number of expressed miRNAs (blue color) and their putative target TF mRNAs (Red colour) in ESC, HC11(N), (P), and (PRL) cell stages, and the number of miRNAs and TF mRNAs either up- or down-regulated between ESC-HC11 (N), N-P, and P-PRL cell stages. Putative TF mediated gene pathways controlled by their respective stage-specific miRNAs in ESC (B), HC11-N (C), P (D), and PRL (E). A representative miRNA-mRNA interactome of affected pathways, such as the Thyroid hormone signalling pathway (B’), TGF-beta signalling pathway (C’), FoxO signalling pathway (D’), and Wnt signalling pathway (E’), is depicted for each category.

### miRNA-ER mRNAs interactome analysis

In a similar manner as TFs, even the interaction maps for Epigenetic regulators (ERs) and miRNAs have been analysed. We found a total of 230 miRNAs that are targeting 434 ERs in ESC (Fig. 7A). In HC11(N), 87 miRNAs targeted 263 ERs. These miRNAs: ER interactions are involved in controlling thyroid cancer, cell cycle, Bacterial invasion of epithelial cells, Pathways in cancer, Wnt signaling pathway, etc. (Fig. 7B). The pathways that came up in HC11(P) are Bacterial invasion of epithelial cells, cell cycle, FoxO signaling pathway, Colorectal cancer, Wnt signaling pathway, Adherens junction, Notch signaling pathway, etc. (Fig. 7C). Similarly, in HC11(P) 96 miRNAs are controlling a total of 274 ERs and the pathways that are involved are Bacterial invasion of epithelial cells, Thyroid hormone signaling pathway, cell cycle, FoxO signaling pathway, Pathways in cancer, Notch signaling pathway, Endometrial cancer, etc. (Fig. 7D). Whereas 234 ERs in the case of HC11(PRL) are targeted by 78 miRNAs which regulate pathways like the Thyroid hormone signaling pathway, Chronic myeloid leukemia, Cell cycle, Bacterial invasion in epithelial cells, non-homologous end joining, etc. (Fig. 7E). Further analysing the differential ERs along with miRNAs provided more specific stage-specific interactions. There were 70 upregulated miRNAs targeting 129 ERs and 40 downregulated miRNAs targeting 69 ERs in HC11(N) vs ESC. Similarly, in HC11(P) vs HC11(N) there were five up miRNAs targeting 16 down ERs and 12 down miRNAs targeting 23 ERs. However, we ended up with only one up miRNA targeting one down ER gene in the case of HC11(PRL) vs HC11(P) (Fig. 7A).

**Fig. 7.**
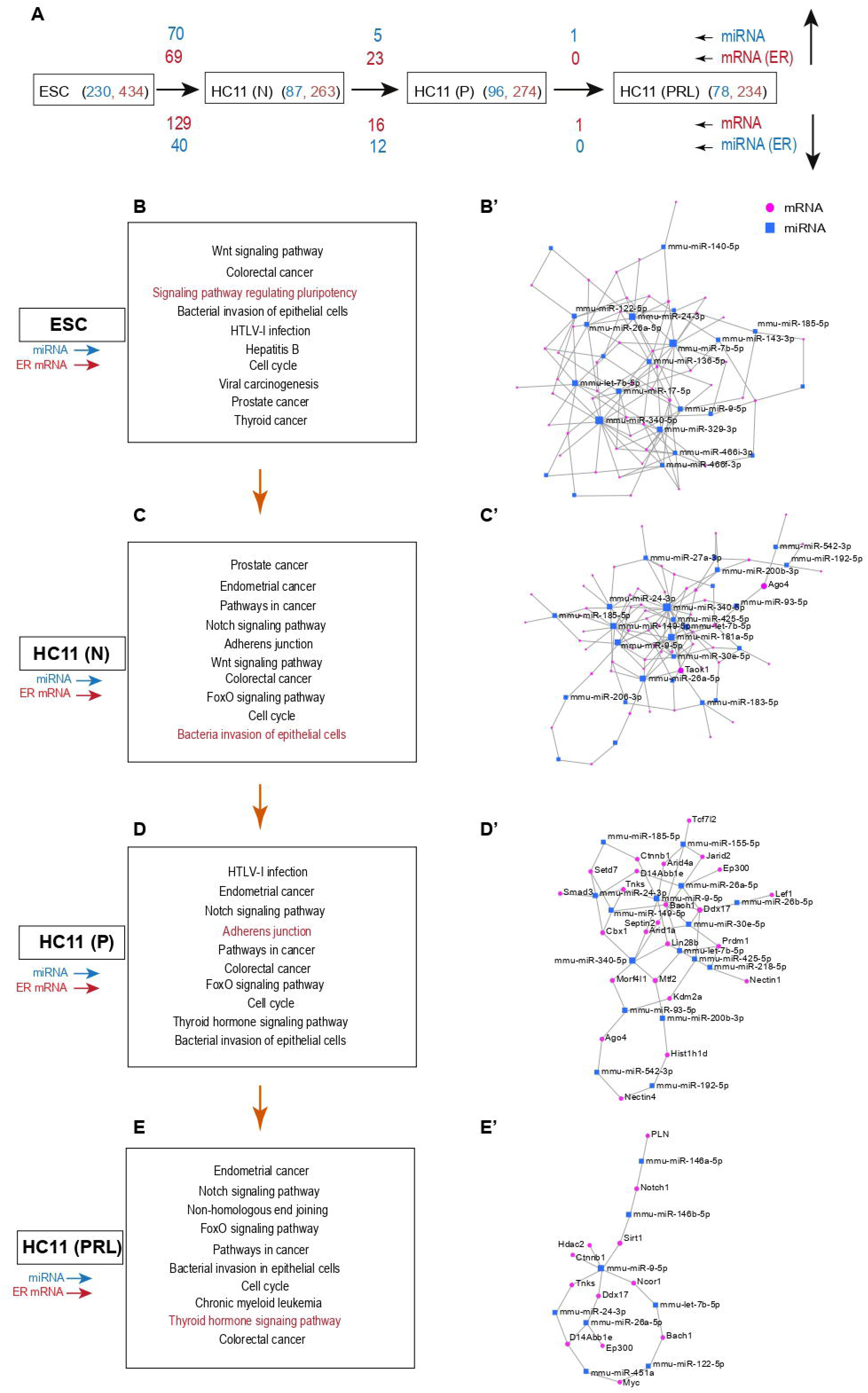
Post-transcriptional control of stage-specific epigenetic regulator (ER) mRNAs and its specific pathways by miRNAs in ESC, HC11 (N), (P) and (PRL) cell stages. (A) Flow chart representing the total number of expressed miRNAs (blue color) and their putative target ER-mediated mRNAs (Red colour) in ESC, HC11(N), (P), and (PRL) cell stages, and the number of miRNAs and ER-mediated mRNAs either upregulated or downregulated between ESC-HC11 (N), N-P, and P-PRL cell stages. Putative ER-mediated gene pathways controlled by their respective stage-specific miRNAs in ESC (B), HC11-N (C), P (D), and PRL (E). A representative miRNA-mRNA interactome of affected pathways, such as signalling pathway regulating pluripotency (B’), Bacterial invasion of epithelial cell pathway (C’), Adherens junction pathway (D’), and Thyroid hormone signaling pathway (E’). ER: Epigenetic regulators.

### Dynamic expression of microRNA genes and their relationship with genomic location during lactogenic differentiation

Cell differentiation is accompanied by the relocalization of genes between heterochromatin (LADs) and euchromatin (inter LADs and fLADs) domains. Keeping this view in mind, we annotated the location of expressed and differentially expressed miRNA genes to the constitutive LADs and constitutive inter LAD regions. This data suggests (Sup. Fig. 6 A, B & C) that under HC11-P conditions multiple miRNA genes present in cLAD regions were selectively upregulated, suggesting a possible role of GR signalling in mediating activation of LAD-embedded miRNA genes.

## Discussion

Embryonic stem cell state represents a hyperdynamic pluripotent state of development. During ESC maintenance, a delicate balance between self-renewal and differentiation is necessary for the proper maintenance and differentiation of ESCs. Multiple mRNA transcripts are transcribed but controlled by miRNAs to prevent ESC differentiation. In line with this assumption, we found that multiple miRNAs are abundant in the ESC state, and our target prediction strategy has identified that comparing MEC with the ground level of pluripotency (ESC) helped us unlock many important miRNA-mRNA interactions that are vital for maintenance and development. ESCs are pluripotent, but MECs are biopotent in nature (Ref). We observed nearly triple the number of expressed miRNAs in ESC as compared to HC11(N), and nearly 249 unique miRNAs were expressed in ESC. The pathways between ESC and MSC were compared in terms of miRNA interactions. The pathways mainly altered are engaged in regulating proliferation, such as the Signaling pathways that regulate the pluripotency of stem cells, Pathways in cancer, the Hippo signaling pathway, the PI3K-Akt signaling pathway, and developmental differentiation pathways, including Focal adhesion, Apoptosis, the AMPK signaling pathway, the Rap1 signaling pathway, and ECM receptor interaction.

Our previous study (Sornapudi et al., 2018) demonstrated that lactogenesis is accompanied by cell cycle arrest; therefore, the dramatic alterations in miRNA expression observed during the different stages of lactogenic differentiation of HC11 also occurred in the absence of cell cycle progression. Interestingly, it has been observed that no miRNAs were downregulated with our cutoff during the MEC (P) to MEC (Prl) transition. However, most miRNAs maintain similar expression levels during MEC(Prl) stages. These observations indicate that significant changes in miRNA transcript levels were mediated under the condition of ECM and GR signalling and were maintained even after the addition of the MEC(Prl) hormone.

To understand the functional state of MECs during their lactogenic differentiation, Kyoto Encyclopedia of Genes and Genomes (KEGG) pathways were derived by integrative mapping of miRNAs and their targeted mRNAs (Fig. 3-7). It was noted that a few miRNAs, such as mmu-mir-340-5p, 19b-3p, 30e-5p, 185-5p, 292a-5p, 203-3p, let-7e-5p, and let-7c-5p, were commonly involved in the majority of the regulatory pathways. Among these, upregulated pathways upon glucocorticoid treatment included Focal adhesion, which is essential for cell binding to the extracellular matrix and has been shown to support ductal branching and morphogenesis during mammary gland development (Paavolinen et al., 2021; Biswas et al., 2022). Here, we found the upregulation of genes Msi2, Col27a1, Col3a1, Cav1, Pik3r1 and Spp1 related to FoxO signaling those were targeted by mmu-mir-340-5p and let-7e-5p in HC11(N) (Supp. Fig. 2A). The upregulation of the FoxO signalling was also observed in HC11(P) vs HC11(N), which plays a critical role in the maintenance of mammary stem cell homeostasis (Sreekumar et al., 2017). Our integrative analysis has shown that the upregulation of mRNAs related to FoxO signalling was correlated with the downregulation of miRNAs such as mmu-mir-19b-3p, 340-5p, and 30e-5p upon glucocorticoid treatment, which is predicted to control Foxo3, Homer2, Gpcpd1, Sfrs18, and Pik3r1 (Supp. Fig. 2B).

The ECM receptor pathway regulates mammary epithelial cell polarization and contributes to milk protein expression, which is upregulated during the developmental transition from MEC(N) to MEC(P) stages, following the initiation of differentiation, transformation, and cell migration in response to hormonal treatment. In the event of any malfunction, the highly controlled ECM receptor pathway during mammary cell differentiation could lead to the advancement of cancer (Zhu et al., 2014) (Supp. Fig. 2C). Another significant pathway that was noticeably upregulated when glucocorticoids were added is the prolactin signalling pathway. It causes alveolar proliferation and terminal differentiation (Hennighausen et al., 2001). On the other hand, it also activates the Jak-STAT pathway. The Jak-STAT pathway modulates the temporal expression of many transcripts required for lactogenic differentiation (Lavnilovitch et al., 2002). Prolactin signalling pathway upregulation was correlated with the downregulation of mmu-mir-340-5p and mmu-mir-19b-3p and Upregulation of Pik3r1, Sfrs18, and Foxo3 (Supp. Fig. 2D). In the end, a fully functional mammary gland requires the Oxytocin signalling pathway to eject milk by causing contraction of myoepithelial cells (Lollivier et al., 2006) which was potentially modulated by mmu-mir-30e-5p, mmu-mir-19b-3p and mmu-mir-340-5p.

In the case of HC11(PRL) vs HC11(P), the commonly involved miRNAs were Mmu-mir-340-5p, let-7e-5p, 122-5p, 19b-3p, 16-5p, 30e-5p, 30a-5p, 30d-5p and let-7c-5p. Among them, mTOR signalling showed significant Upregulation upon Prolactin treatment and is essential for mammary epithelial cell proliferation and differentiation. In a previous study, Jankiewicz et al. (2006) observed a reduction in milk protein expression in cultured cells following the inhibition of the mTOR pathway. Here, the mTOR signalling pathway was accompanied by the downregulation of mmu-mir-30e-5p and 340-5p, and an increase in the expression of Vegfa, Slc7a1, and Ddit4 (Supp. Fig. 3C).

The PI3K/Akt pathway is very much necessary for the proper functioning of the mammary gland. Overexpression of this had an impact on delaying the involution process (Wickenden et al., 2010). In our study, it was found to be regulated by mmu-mir-340-5p, 122-5p, and 30e-5p by targeting Myc, Vegfa, Slc7a1, and Ddit4 (Supp. Fig. 3B). The upregulation of the MAPK signalling pathway was shown to be vital for cell survival as its inhibition was shown to promote apoptosis (Healy et al., 2000). MAPK signalling pathway-specific mRNAs in MEC(P) and MEC(Prl) stages were predicted to be regulated by downregulation of miRNAs such as mmu-mir-19b-3p, 340-5p, and 122-5p. Those that promote upregulation of Myc, Slc7a1, Sfrs18, and Gadd45a (Supp. Fig. 3C). The Wnt signalling pathway is critical for mammary gland growth, differentiation, and involution (Turashvill et al., 2006). In the MEC (Prl) stage, mmu-mir-122-5p and 292a-3p were identified to be involved in the regulation of Wnt signalling-specific mRNAs. Here, mmu-mir-292a-3p targets the Dkk-1 gene, which is responsible for initiating apoptosis by inhibiting the Wnt signaling pathway. That is how mammary cells stabilize Wnt signaling, which is essential during the Prolactin state. Estrogen signaling plays a crucial role in the development of the mammary gland during the pubertal stage by facilitating ductal morphogenesis (LaMarca et al., 2007), as predicted to be regulated by mmu-miR-30e-5p at the MEC (PRL) stage.

Furthermore, downregulated miRNAs were analyzed explicitly in conjunction with filtered upregulated TFs, revealing the upregulation of particular TF targets of miRNAs, rather than a complex transcriptomic network. Interaction maps of HC11(N) vs ESC consisted of a relatively large number of miRNAs: TF interactions, which were involved in regulating pathways related to bacterial invasion of epithelial cells, Lysin degradation, and Metabolic pathways. The list of downregulated miRNAs, mainly associated with upregulated TFs during the HC11(N) to HC11(P) transition, included mmu-mir-340-5p, 19b-3p, 30e-5p, and 185-5p. Downregulation of mmu-miR-19b-3p led to the Upregulation of Tef, which is involved in the Hippo signalling pathway and is found to be essential during pregnancy (Chen et al., 2014). Zhx3 was predicted as a target of both mmu-mir-19b-3p and mmu-miR-30e-5p in the Primed state. In contrast, Nfat5 is a target of both mmu-miR-30e-5p and mmu-miR-185-5p. Likewise, a vice versa analysis of upregulated miRNA and downregulated TFs in HC11(N) vs ESC identified mmu-mir-let-7b-5p, 181a-5p, 30e-5p, 122-5p, and 340-5p, which control transcriptional misregulation in cancer, signaling pathway regulation of pluripotency, and pathways in cancer (Fig. 6).

Similarly, the targets of miRNAs among epigenetic regulators were identified. The comparison of HC11(N) with ground ESC by considering down mi and up ERs provided the following list of miRNAs, mmu-mir-7b-5p, 129-5p, 466f-3p, 495-3p, 329-3p, 410-3p, etc., with more hits of ERs, and the targeting pathways under these ERs are Adherens junction, FoxO signaling pathway, Hippo signaling pathway, Wnt signaling pathway, etc. A total of 12 miRNAs (mmu-mir-340-5p, 19b-3p, 30e-5p, 122-5p, 16-5p, 292a-5p, 203-3p, let-7c-5p, let-7e-5p, 185-5p, 146a-5p, and 291a-3p) were identified in our dataset as specifically targeting ERs during the transition towards the HC11(P) state from HC11(N). However, considering the up-and-down miRNAs in the primed state, identified five significant miRNAs. Here, Mmu-mir-155-5p is predicted to target Myb, which is essential to control tumorigenesis during mammary gland development (Miao et al., 2011). Furthermore, Sox11 is an embryonic mammary marker that remains silent during the postnatal development of the mammary gland (Umeh-Garcia et al., 2020), which is regulated by mmu-miR-204-5p.

Interestingly, there are three miRNAs among them, mmu-mir-340-5p, 30e-5p, and 122-5p, whose expressions were significantly upregulated in HC11(N) in comparison to ground ESC. However, the expressions were differentially downregulated towards primed differentiation in addition to glucocorticoid. These miRNAs occupied the central hub in the interaction maps of up miRNA: down mRNA of HC11(N)_ESC and down miRNA: up mRNA of HC11(P)_HC11(N). They also have significant targets in our datasets that play vital roles in mammary gland development.

## Conclusion

The study focuses on the development of the mammary gland by performing high-throughput miRNA sequencing of Embryonic Stem Cells (ESC) and mouse mammary epithelial stem-like cells (HC11) during lactogenic differentiation, revealing many differentially expressed miRNAs. Here, we have shown the dynamic expression of thousands of miRNAs in a stage-specific manner. This study utilizes high-throughput next-generation sequencing to investigate the potential role of miRNAs in post-transcriptional gene expression control. The study also analyzed various miRNA-mRNA interactome maps of up- and downregulated genes, identifying significant interactions essential for mammary gland differentiation and development.

| Sl No. | MiRNA | Role in mammary gland | References |
| --- | --- | --- | --- |
| 1 | MiR-122-5p | Promotes breast cancer metastasis | Lv et al., 2025 |
| 2 | Let-7a-1 | Downregulates breast cancer cell invasion and migration | Thammaiah et al., 2016 |
| 3 | Let-7c-2 | Involved in progression of breast tumors | Bozgeyik, 2020 |
| 4 | MiR-19b | Tumor progression in breast cancer | Li et al., 2018 |
| 5 | MiR-155-5p | Breast cancer development and progression | Pasculli et al., 2020 |
| 6 | MiR-1a-3p | Inhibits migration and promote apoptosis in breast cancer | Dai et al., 2023 |
| 7 | MiR-149-5p | Role in tumor suppressor | Fard et al., 2021 |

## Materials and Methods

### 1. Cell culturing and lactogenic differentiation of HC11 MECs

HC11 MECs were cultured for lactogenic differentiation as described by Sonrnapudi et al. (2018). Briefly, two independents freshly thawed HC11 cells (Passage number 4) were cultured and expanded in three T25 flasks (Corning #420639) in 10% complete media (DMEM supplemented with 10% FBS (A5256701) and 1X Antibiotic Antimitotic), Insulin (5μg/ml; Sigma # 16634) and EGF (20ng/ml; Sigma # E4127) at 37°C, 5% CO2 in incubator, until the cells reached confluency. After reaching confluency, cells from one of the flasks were harvested and treated as HC11 (N) stage of cells. The remaining media from the other two flasks were replaced with 5% complete medium (DMEM supplemented with 5% FBS and 1X Antibiotic-Antimycotic), Hydrocortisone (1 μg/ml; Sigma #H4001), and Insulin (5 μg/ml), and then the cells were cultured for 48 hours under the same conditions. After 48 hours, cells from one of the flasks were harvested and treated with HC11 (P) stage of cells. The medium in the third flask was replaced with 5% complete medium supplemented with prolactin hormone (NIH # NIDDKoPRL-21) (5 μg/ml) and cultured for up to 72 hours. After 72 hours, cells were harvested and treated as HC11 (PRL) stage of cells. Preservation: HC11 cells were preserved by growing them under Insulin and EGF to 50-60% confluency. After trypsinization (1X, for 5-10 min at 37°C), the cells were resuspended in DMEM media containing 7% DMSO and 20% FBS. They were then placed in a −20°C freezer for 1 hour, followed by −80°C for 2 hours, and finally into liquid nitrogen for long-term storage.

### 2. Cell culturing of murine R1 Embryonic Stem cells

R1 ESCs were thawed in a 37°C water bath for 5 minutes, mixed with 10 mL of 10% ESC medium, and then collected by centrifugation at 800 RPM for 5 minutes. The cell pellet was resuspended in the ESC medium and placed on a previously treated 0.1% gelatin-coated Petri plate containing a mitomycin-C-inactivated monolayer of dormant mouse embryonic fibroblast feeder cells (Bibel et al., 2007). These cells were cultured to ∼80% confluency at 37°C with 7% CO2 in the presence of 10% FBS, LIF, and 2i (MEK 0.5 μg/ml: GSK: 1.5 μg/ml) medium. Later, ESCs were selected over fibroblasts using a trypsinization method; a 10 mL mixture of fibroblasts and ESCs was added to a 100 mm petri dish to enrich the ESCs from the fibroblasts selectively. Since fibroblasts adhere to the surface more quickly than ESC, they were incubated for 30 minutes at 37°C and 7% CO2 to allow for settling. Afterward, a complete 10 ml medium was collected slowly and transferred to another gelatin-coated petri dish with 10% ESC medium. Three cycles of the above selective enrichment process were performed to obtain a nearly 90% pure population of ESCs, free from fibroblast cells. These ESCs were harvested for the isolation of miRNAs, preparation of a miRNA sequencing library, and high-throughput Illumina sequencing.

### 3. Isolation of mRNA and microRNAs from ESC, HC11 MECs (N), (P), and (PRL) stages

MicroRNAs from HC11 cells undergoing lactogenic differentiation (N, P, and PRL) and ESC were isolated by using miRVana miRNA isolation Kit (Invitrogen# AM1560) as per the recommendations with minor modifications. Briefly, 300μl of lysis buffer (Provided in the miRNA isolation Kit) was added directly to 3-5 million cells (cells from one T25/30-60mm Petri plate) at room temperature (RT). 30 μL of miRNA homogenate additive was added and incubated for an additional 10 minutes on ice. Then, 300 μL acid-phenol: chloroform in a 1:1 ratio was added, mixed by vortexing, and centrifuged at 10,000rpm for 5min at room temperature. The aqueous phase was separated, and 100 μL of absolute ethanol (one-third of the total volume of the aqueous phase) was added and mixed thoroughly. It was mixed and passed through the spin column-1 by centrifugation at 10,000RPM for 1min at RT. Spin column-1 was kept on a 2ml tube for mRNA extraction. 266μL 100% ethanol (2/3th of the total flow through) was added to the flow through. The eluent (which contains miRNAs) was mixed and passed through column-2 by centrifugation at 10,000RPM for 1min at RT. Both column1 and column2 were washed with 700 μL of miRNA Wash Solution-1, followed by 2 times 500μL Wash Solution 2/3 by centrifugation at 10,000RPM for 1min at RT. Columns were wet-dried by centrifugation at 10K RPM for 2min at RT. 50μl and 30μl nuclease-free water were added into column-1 and column-2 respectively. Columns were incubated for 1min at RT and centrifuged at 10,000 RPM for 1min. Isolated miRNA and mRNA concentration was measured either by using a spectrophotometer (nanodrop) or Qubit RNA HS test kit (Thermo Fisher#Q32855) using Qubit Fluorometer 4.0.

### 4. MicroRNA Library Preparation and Sequencing

Fluorimeter, the Qubit RNA HS Kit (Invitrogen # Q32852) was used to measure the RNA concentration. All samples were analyzed using an Agilent 2100 Bioanalyzer to assess the integrity of the isolated RNA. The NEBNext Small RNA Library Prep Set for Illumina (NEB# E7330L) was used to generate miRNA libraries. Additionally, an Illumina HiSeq 2500 platform was used for Illumina single-end sequencing of miRNA-seq (1X50bp).

### 5. Preparation of cDNA

MicroRNA cDNA was produced with the use of the miScript II RT kit (Qiagen #218161). A thermocycler was used to incubate 1 μg miRNA, 4 μl HiSpec buffer, 2 μl nucleotide, 1 μl reverse transcriptase enzyme, and nuclease-free water up to 20 μl for 60 minutes at 37 °C and then for 5 minutes at 95 °C.

### 6. Primer Designing and RT-PCR

The exact mature miRNA sequence was considered as miRNA PCR forward primers. Only the nucleotide U was changed to T for cDNA primers. A universal reverse primer was used from the miScript SYBR Green PCR kit (Qiagen #218073). miRNA cDNA was diluted with NF water up to 20ng/μl concentration. The total reaction volume of each well was 10μl (5μl 2X miScript SYBR Green PCR kit, 1μl miRNA cDNA template, 1μl of 5μm primer mix, and 3μl NF water). RT-PCR was performed using a CFX96 Touch Real-Time PCR (Bio-Rad) and the conditions were set according to the miScript SYBR Green PCR kit (Qiagen #218073). The *Rnu6* gene was used as a housekeeping gene. All the data were analysed by using the −2^ΔΔ^CT method (Livak & Schmittgen, 2001). The *Rnu6* gene and HC11 (N) sample were considered as the control. Bar graphs were generated with one-way ANOVA by using Graphpad PRISM software. (Supplementary file showing microRNA gene name and primer sequences used for RT PCR analysis).

### 7. Bioinformatic analysis of miRNA seq data

**(a) Generation of normalized count:** 20 million single-end reads were obtained for each sample (Supplementary table showing stage, replicates, raw sequences, aligned reads, mapped score etc). Sequenced reads were initially processed for quality control with FastQC (www.bioinformatics.babraham.ac.uk/projects/fastqc/). Adapter sequences were removed with Cutadapt, and Spearman’s correlation was analysed between replicates by using R. High-quality sequence reads were mapped to the mouse reference genome (mm10) by using the Bowtie alignment tool (bowtie-bio.sourceforge.net/manual.shtml). Mapped reads were processed through miRDeep2 (Friedlander et al., 2011) to predict normalized count for both known and novel miRNAs. Datasets were imported to Excel (Supplementary table showing counts against microRNAs from ESC, N, P, and PRL stages). Normalized count values above 10 were considered as expressed microRNAs in a particular state, i.e., ESC, HC11 (N), (P), and (PRL) stages.
**(b) Analysis of differentially expressed miRNAs:** Output file (Text file format) of miRDeep2 was processed further with DESeq2 to generate differentially expressed miRNAs between ESC-HC11 (N), N-P, and P-PRL stages. From the differentially expressed miRNAs list, Log2 fold change above 1 with P value < 0.01 were considered as upregulated genes and Log2 fold change below −1 with P value < 0.01 as downregulated genes.
**(c) Analysis of uniquely expressed miRNA genes in ESC, HC11 (N), (P), and (PRL) states:** Uniquely expressed miRNA list was extracted from the Venn diagram for all the states separately (Fig. 1C) and assigned with its respective normalized count. Among them, the top 10 miRNAs with the highest value of normalized count were considered for the heatmap (Fig. 2B)
**(d) Analysis of highly expressed miRNA genes in ESC, HC11 (N), (P), and (PRL) states:** At first, common 97 miRNAs among ESC, HC11(N), (P), and (PRL) were eliminated from the main list. The top 10 miRNAs with highest value of normalized count was considered for highly expressed miRNAs in respective states (FIg. 2A).
**(e) Analysis of highly upregulated miRNA genes between cellular states:** For highly upregulated miRNAs, Log2 fold change value above or equal to 1 of genes between stages were considered significant. Among them, the top 10 miRNAs with the highest log2 fold values were considered for the heatmap (Fig. 2C).
**(f) Analysis of highly down-regulated miRNA genes between cellular states:** For highly downregulated miRNAs, Log2 fold change values below or equal to −1 were considered significant. Among these, the 10 miRNAs with the lowest log2 fold values were considered for the heatmap (Fig. 2D).
**(g) Analysis of miRNAs targeting transcription factor (TFs) mRNAs in ESC, HC11 (N) (P) and (PRL) states:** To analyse the list of TFs that are targeted by stage-specific miRNAs, targeted gene list was filtered out with stage specific TFs as per Sornapudi et al., 2018. Afterwards, pathways have been derived by considering these targeted TFs.
**(h) Analysis of miRNAs targeting epigenetic regulators (ERs) mRNAs in ESC, HC11 (N), (P), and (PRL) stages:** To analyze the list of ERs, the targeted gene list was filtered out with stage specific ERs as per Sornapudi et al., 2018, and pathways were also derived similarly.

### 8. Integrative miRNA and mRNA global interactome analysis

**(a)** Integrative analysis of mRNA and microRNA Interactome analysis in ESC, HC11 (N), (P), and (PRL) stages. mRNA-seq data from previously published datasets (GSE107419) from ESC, HC11 (N), (P), and (PRL) stages were considered for integration with the respective stage-specific microRNA-seq data generated in this study. (i) ESC mRNA vs miRNAs, (ii) HC11 (N) mRNAs vs miRNAs, (iii) HC11 (P) mRNAs vs miRNAs, (iv) HC11 (PRL) mRNA vs miRNAs.
**(b) Down-regulated miRNA-up-regulated mRNA genes interactome analysis during cell state transition:** Differential upregulated interaction maps were generated between HC11(N)_ESC by considering upregulated mRNAs with Log2 fold change ≥1 and downregulated miRNAs with Log2 fold change ≤-1. In the case of HC11(P)_HC11(N), downregulated miRNAs of HC11(P)_HC11(N) were only considered because no network was generated, while upregulated mRNAs of HC11(P)_HC11(N) were batch filtered from the targeted network list. For the interaction map of HC11(Prl)_HC11(P), upregulated mRNAs with Log2 fold change ≥1 of HC11(Prl)_HC11(P) and downregulated miRNAs with Log2 fold change ≤-1 of HC11(P)_HC11(N) were considered.
**(c) Upregulated miRNA-downregulated mRNA genes interactome analysis during cell state transition:** Differential downregulated interaction maps were generated between HC11(N)_ESC, HC11(P)_HC11(N), and HC11(Prl)_HC11(P) by considering downregulated mRNAs with Log2 fold change ≤-1 and upregulated miRNAs with Log2 fold change ≥1.
**(d) Upregulated miRNA-mRNA genes interactome analysis during cell state transitions:** Differential upregulated miRNA-mRNA interaction maps were generated between HC11(N)_ESC, HC11(P)_HC11(N), and HC11(Prl)_HC11(P) by considering upregulated mRNAs with Log2 fold change ≥1 and upregulated miRNAs with Log2 fold change ≥1.
**(e) Down-regulated miRNA-mRNA genes interactome analysis during cell state transitions:** Differential upregulated miRNA-mRNA interaction maps were generated between HC11(N)_ESC, HC11(P)_HC11(N), and HC11(Prl)_HC11(P) by considering downregulated mRNAs with Log2 fold change ≤-1 and downregulated miRNAs with Log2 fold change ≤-1.

### 9. Pathway analysis

KEGG pathway analysis and Gene Ontology (GO) study were conducted for the above list of miRNA and mRNA combinations (from sections 7 and 8 of the method) in miRNet, considering a P-value below 0.1 for both KEGG pathway and GO analysis. Furthermore, pathway-specific interaction maps were extracted from the leading network to provide a detailed overview of the interactions within each pathway.

### 10. Analysis of genomic locations

Analysis of genomic locations of miRNAs in cLAD, ciLAD, fLAD and No-LAD regions: The boundaries of cLADs, ciLADs, and fLADs were downloaded from GSE17051 (Peric-Hupkes et al., 2010). The genomic regions which were not annotated for LADs were considered as No-LAD regions. These mm9 coordinates were converted to mm10 using the LiftOver tool in UCSC genome browser. List of miRNA genes were retrieved from mouse GRCm38/mm10 assembly from the UCSC genome browser. The coordinates of conserved-LADs conserved inter-LADs and facultative-regions were retrieved from available datasets (Peric-Hupkes et al., 2010). Based on the coordinates of miRNA genes, we assigned genes to either conserved c/ci/f/No-LADs. All the genes that have a minimal FPKM value were considered a list of genes expressed in ESC, HC11-N, HC11-P and HC11-PRL cell types and differentially expressed miRNA genes between ESC-HC11(N), HC11 (N)-HC11 (P) and HC11 (P)-HC11 (PRL) stages.

## 11. Data Availability

MiRNA sequencing data will be publicly available through the GEO; GSE137388.

## Supporting information

Statistics of miRNA-sequencing datasets.

mRNA-seq and miRNA-seq normalized counts of mRNAs (Up or down-regulated) and miRNAs (Down or up-regulated) between ESC and HC11 (N) stages.

mRNA-seq and miRNA-seq normalized counts of up-regulated mRNAs and down-regulated miRNAs between HC11 (N)-(P) cell stages.

mRNA-seq and miRNA-seq normalized counts of up or down regulated mRNAs and down and up regulated miRNAs between HC11 (N)-(P) cell stages.

mRNA-seq and miRNA-seq normalized counts of up- or down-regulated mRNAs and down and up-regulated miRNAs between HC11 (P)-(PRL) cell stages.

Genomic distribution of expressed and differentially expressed miRNAs during lactogenic differentiation

List of miRNAs and genes with the forward and reverse primers used for the Real time PCR.

Sup. Fig. 1. **Statistics of miRNA-sequencing datasets**. (A) Table showing the number of reads sequenced, aligned reads per replicate of ESC, HC11 (N), (P), and (PRL) samples. (B&C) PCA analysis of biological replicates of ESC, HC11(N), HC11(P), and HC11(PRL) datasets.

Sup. Fig. 2. **mRNA-seq and miRNA-seq normalized counts of mRNAs (Up or down-regulated) and miRNAs (Down or up-regulated) between ESC and HC11 (N) stages**. (A1) mRNA-miRNA interactome of upregulated Apoptosis pathway mRNA genes due to down regulation of miRNAs between ESC-HC11 (N) stages and its corresponding FPKM values of mRNAs (A2) and normalized counts of miRNAs (A3) represented by bar charts.

Sup. Fig.3. **mRNA-seq and miRNA-seq normalized counts of up-regulated mRNAs and down-regulated miRNAs between HC11 (N)-(P) cell stages**. (A1) mRNA-miRNA interactome of upregulated prolactin signalling pathway mRNA genes due to down regulation of miRNAs between HC11 (N)-(P) stages and its corresponding FPKM values of mRNAs (A2) and normalized counts of miRNAs (A3) represented by bar charts. (B1,2&3) for Oxytocin signalling pathway, (C1,2&3) FoxO signalling, (D1,2&3) Focal adhesion, (E1,2&3) ECM receptor interactions.

Sup. Fig.4. **mRNA-seq and miRNA-seq normalized counts of up or down regulated mRNAs and down and up regulated miRNAs between HC11 (N)-(P) cell stages**. (A1) mRNA-miRNA interactome of upregulated prolactin signalling pathway mRNA genes due to downregulation of miRNAs between HC11 (N)-(P) stages and its corresponding FPKM values of mRNAs (A2) and normalized counts of miRNAs (A3) represented by bar charts. (B1,2&3) for the Bacterial invasion pathway, (C1,2&3) downregulated the P53 signalling pathway mRNAs by upregulating miRNAs.

Sup. Fig.5. **mRNA-seq and miRNA-seq normalized counts of up- or down-regulated mRNAs and down and up-regulated miRNAs between HC11 (P)-(PRL) cell stages**. (A1) mRNA-miRNA interactome of upregulated MAPK signalling pathway mRNA genes due to the regulation of miRNAs between HC11 (P)-(PRL) stages and their corresponding FPKM values of mRNAs (A2) and normalized counts of miRNAs (A3) represented by bar charts. (B1,2&3) for the mTOR pathway, (C1,2&3) PI3K-AKT signalling pathway, (D1,2&3) downregulated Wnt signalling pathway mRNAs by upregulated miRNAs.

Sup. Fig.6. **Genomic distribution of expressed and differentially expressed miRNAs during lactogenic differentiation**: (A) Schematic representation of number of expressed and differentially expressed miRNAs coming from c, ci, f and No LAD regions in ESC, HC11-N, P and PRL states (B) Venn diagram showing common and unique miRNA originated from c, ci, f and No LAD regions in ESC, HC11-N, P and PRL states. (C) Bar chart showing the relative number of expressed miRNAs coming from c, ci,f, and No LAD regions in ESC, HC11-N, P, and PRL states

Table1. List of genes with the forward and reverse primers used for the Real time PCR.

Table2. List of MiRNAs with the forward primers used for the Real time PCR.

## References

1. Accili, D., & Arden, K. C. (2004). FoxOs at the crossroads of cellular metabolism, differentiation, and transformation. Cell, 117(4), 421–426. 10.1016/S0092-8674(04)00452-0

2. Bartel, D. P. (2018). Metazoan microRNAs. Cell, 173(1), 20–51. 10.1016/j.cell.2018.03.006

3. Bissels, U., et al. (2012). RNA sequencing of murine mammary epithelial stem-like cells (HC11) undergoing lactogenic differentiation and its comparison with embryonic stem cells. BMC Research Notes, 11, 241. 10.1186/s13104-018-3351-4

4. Biswas, T., Guo, J., & Ewald, A. J. (2022). Extracellular matrix and focal adhesion signaling in mammary gland morphogenesis. Journal of Mammary Gland Biology and Neoplasia, 27(1), 1–15. 10.1007/s10911-021-09500-1

5. Cardoso-Moreira, M., Halbert, J., Valloton, D., Velten, B., et al. (2019). Gene expression across mammalian organ development. Nature, 571(7766), 505–509. 10.1038/s41586-019-1338-5

6. Chang, L., Zhou, G., Soufan, O., & Xia, J. (2020). miRNet 2.0: Network-based visual analytics for miRNA functional analysis and systems biology. Nucleic Acids Research, 48(W1), W244–W251. 10.1093/nar/gkaa467

7. Chen, Q., Zhang, N., Xie, R., Wang, W., Cai, J., Choi, K. S., David, K. K., Huang, B., Yabuta, N., Nojima, H., & Pan, D. (2014). Homeostatic control of Hippo signaling activity revealed by an endogenous activating mutation in YAP. Genes & Development, 28(20), 2221–2234. 10.1101/gad.246488.114

8. Confuorti, C., Jaramillo, M., & Plante, I. (2024). Hormonal regulation of miRNAs during mammary gland development. Biology Open, 13(6), bio060308. 10.1242/bio.060308

9. Davidson, E. H. (2010). Emerging properties of animal gene regulatory networks. Nature, 468(7326), 911–920. 10.1038/nature09645

10. Faherty, S., Fitzgerald, A., Keohan, M., & Quinlan, L. R. (2007). Self-renewal and differentiation of mouse embryonic stem cells as measured by Oct4 expression: The role of the cAMP/PKA pathway. In Vitro Cellular & Developmental Biology – Animal, 43(1), 37–47. 10.1007/s11626-006-9001-5

11. Healy, A. M., et al. (2000). The MAPK pathway and apoptosis in epithelial cells. Molecular Biology of the Cell, 11(8), 2563–2574.

12. Hennighausen, L., Robinson, G. W., Wagner, K. U., & Liu, X. (2001). Prolactin signaling in mammary gland development. Journal of Biological Chemistry, 276(51), 47071–47074. 10.1074/jbc.R100065200

13. Jankiewicz, M., Groner, B., & Desrivieres, S. (2006). Inhibition of mTOR pathway reduces milk protein synthesis in mammary epithelial cells. Molecular Endocrinology, 20(8), 1940–1952.

14. Kanehisa, M., & Goto, S. (2000). KEGG: Kyoto Encyclopedia of Genes and Genomes. Nucleic Acids Research, 28(1), 27–30. 10.1093/nar/28.1.27

15. LaMarca, H. L., Rosen, J. M., & Miner, J. H. (2007). Hormonal regulation of mammary gland development. Endocrinology, 148(7), 3003–3011.

16. Laplante, M., & Sabatini, D. M. (2012). mTOR signaling in growth control and disease. Cell, 149(2), 274–293. 10.1016/j.cell.2012.03.017

17. Lavnilovitch, A., et al. (2002). STAT5 signaling in lactogenic differentiation. Molecular Endocrinology, 16(4), 774–786.

18. Levine, M., & Tjian, R. (2003). Transcription regulation and animal diversity. Nature, 424(6945), 147–151. 10.1038/nature01763

19. Li, Y., et al. (2009). Most mammalian mRNAs are conserved targets of microRNAs. Genome Research, 19(1), 92–105. 10.1101/gr.082701.108

20. Lollivier, V., et al. (2006). Oxytocin signaling and milk ejection. Journal of Dairy Science, 89(1), 13–25.

21. Malizia, A. P., & Wang, D.-Z. (2011). miRNA in cardiomyocyte development. Wiley Interdisciplinary Reviews: Systems Biology and Medicine, 3(2), 183–190. 10.1002/wsbm.111

22. Manning, B. D., & Toker, A. (2017). AKT/PKB signaling: Navigating the network. Cell, 169(3), 381–405. 10.1016/j.cell.2017.04.00

23. McGregor, R. A., & Choi, M. S. (2011). MicroRNAs in the regulation of adipogenesis and obesity. Current Molecular Medicine, 11(4), 304–316. 10.2174/156652411795677990

24. Miao, Y., et al. (2011). Myb controls mammary epithelial proliferation and tumorigenesis. Cancer Research, 71(13), 4431–4442.

25. Paatero, I., Seagroves, T. N., Vaparanta, K., Han, W., Jones, F. E., Johnson, R. S., & Elenius, K. (2014). Hypoxia-inducible factor-1α induces ErbB4 signaling in the differentiating mammary gland. Journal of Biological Chemistry, 289(32), 22459–22469. 10.1074/jbc.M113.533497

26. Paavolinen, L., et al. (2021). Focal adhesion signaling in mammary epithelial morphogenesis. Developmental Biology, 473, 20–33.

27. Peric-Hupkes, D., Meuleman, W., Pagie, L., Bruggeman, S.W., Solovei, I., Brugman, W., Graf, S., Flicek, P., Kerkhoven, R.M., van Lohuizen, M., et al. (2010). Molecular maps of the reorganization of genome-nuclear lamina interactions during differentiation. Mol Cell 38, 603–613. 10.1016/j.molcel.2010.03.016.

28. Perotti, C., et al. (2009). “Characterization of mammary epithelial cell line HC11 using the NIA 15k gene array reveals potential regulators of the undifferentiated and differentiated phenotypes.” Differentiation 78(5): 269–282.

29. Schwanhäusser, B., et al. (2011). Global quantification of mammalian gene expression control. Nature, 473(7347), 337–342. 10.1038/nature10098

30. Sornapudi, T. R., et al. (2018). Transcriptome profiling of HC11 cells reveals lactogenic differentiation dynamics. BMC Research Notes, 11, 241. 10.1186/s13104-018-3351-4

31. Sreekumar, A., et al. (2017). FoxO signaling regulates mammary stem cell homeostasis. Stem Cell Reports, 9(2), 496–509.

32. Stefanie, A.-S., et al. (2009). Characterisation of microRNA expression in post-natal mouse mammary gland development. BMC Genomics, 10, 548. 10.1186/1471-2164-10-548

33. Turashvili, G., et al. (2006). Wnt signaling in mammary gland development and cancer. Journal of Mammary Gland Biology and Neoplasia, 11(3–4), 177–190.

34. Umeh-Garcia, M. C., et al. (2020). Sox11 marks embryonic mammary progenitors. Development, 147(3), dev184960.

35. Wang, W., et al. (2009). “Global expression profiling reveals regulation of CTGF/CCN2 during lactogenic differentiation.” J Cell Commun Signal 3(1): 43–55.

36. Wickenden, J. A., et al. (2010). PI3K/Akt signaling in mammary gland involution. Developmental Biology, 337(2), 219–229.

37. Williams, C., et al. (2009). Gene expression in murine mammary epithelial stem cell-like cells shows similarities to human breast cancer gene expression. Breast Cancer Research, 11(3), R26. 10.1186/bcr2256

38. Wu, D., Thompson, L. U., & Comelli, E. M. (2022). MicroRNAs: A link between mammary gland development and breast cancer. International Journal of Molecular Sciences, 23(24), 15978. 10.3390/ijms232415978

39. Zhu, J., et al. (2014). ECM receptor signaling in mammary epithelial differentiation and cancer. Breast Cancer Research, 16(3), R56.

