## Supplementary figures and images for "MicroRNA-mRNA Gene Regulatory Circuitry Orchestrates the Epigenetic and Transcriptional Reprogramming from Pluripotency to Murine Mammary Epithelial stem-like-Cells lactogenic differentiatio"

### Genomic distribution of expressed and differentially expressed miRNAs during lactogenic differentiation

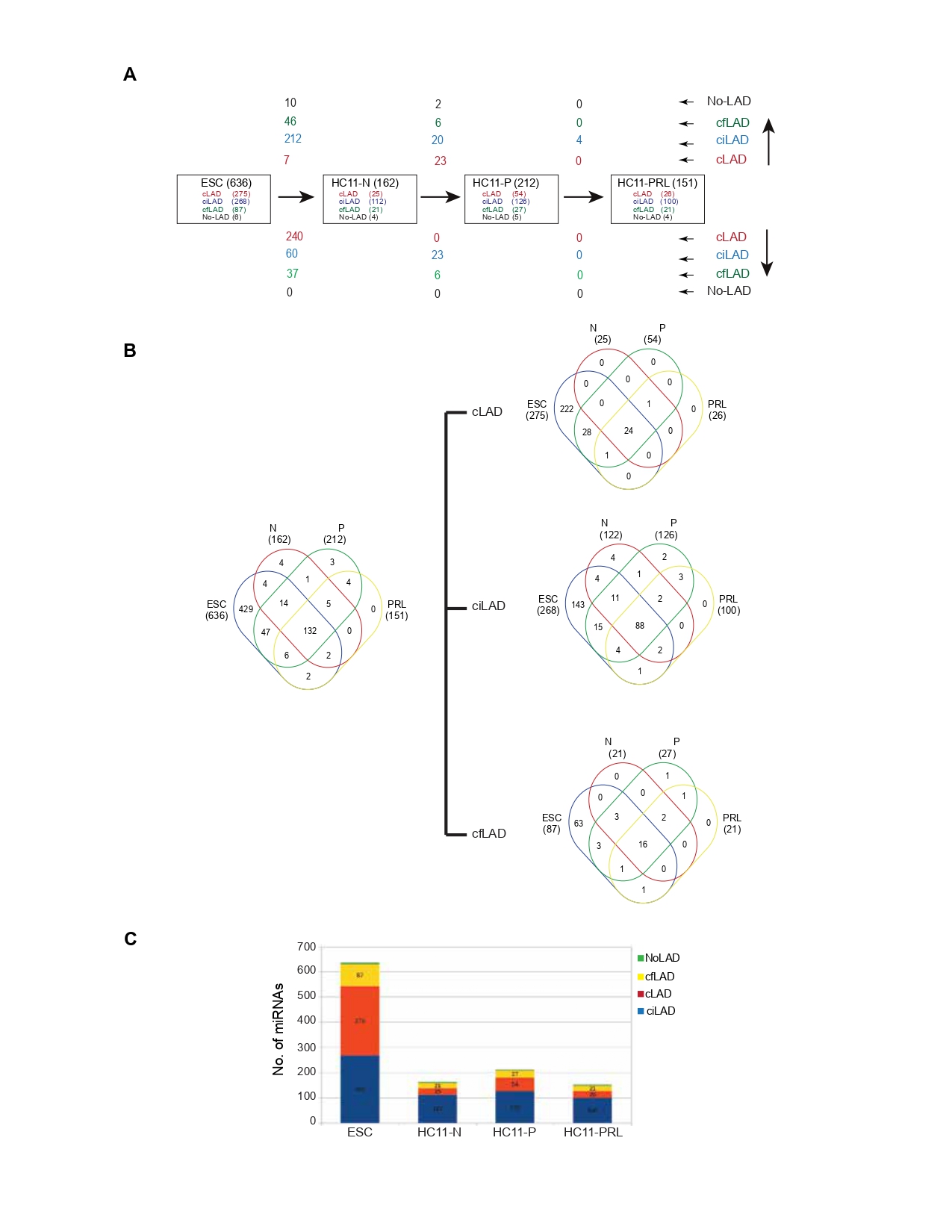

### List of miRNAs and genes with the forward and reverse primers used for the Real time PCR.

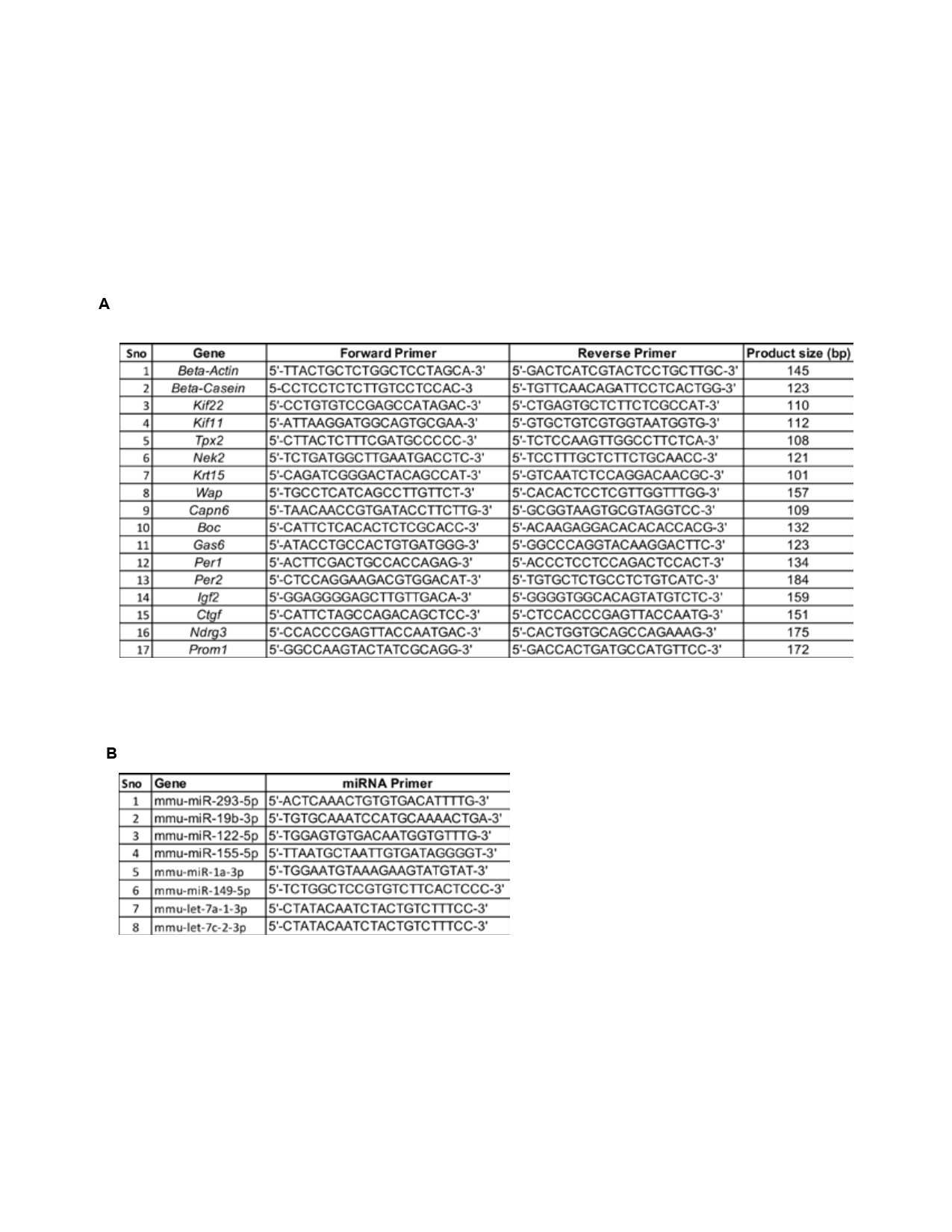

### mRNA-seq and miRNA-seq normalized counts of mRNAs (Up or down-regulated) and miRNAs (Down or up-regulated) between ESC and HC11 (N) stages.

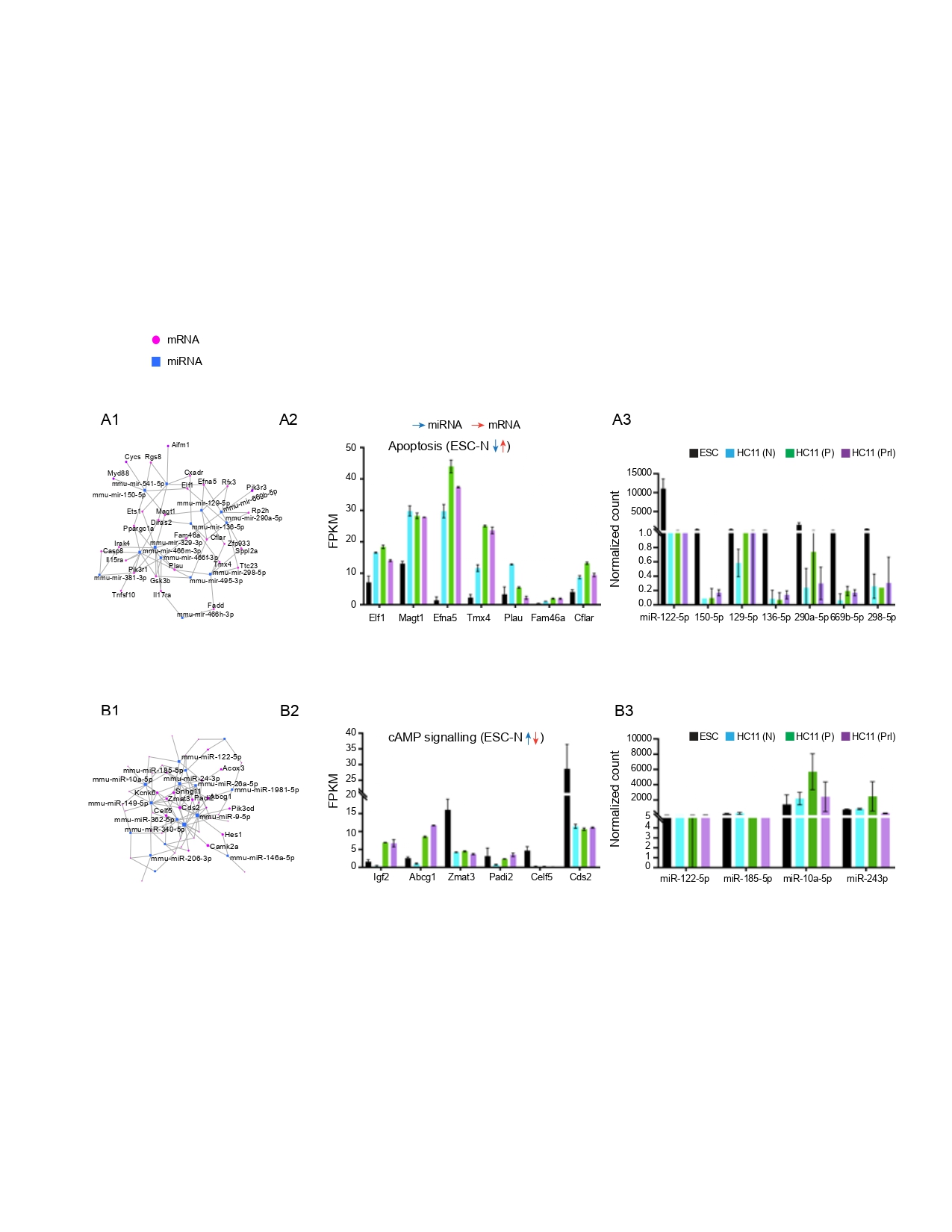

### mRNA-seq and miRNA-seq normalized counts of up or down regulated mRNAs and down and up regulated miRNAs between HC11 (N)-(P) cell stages.

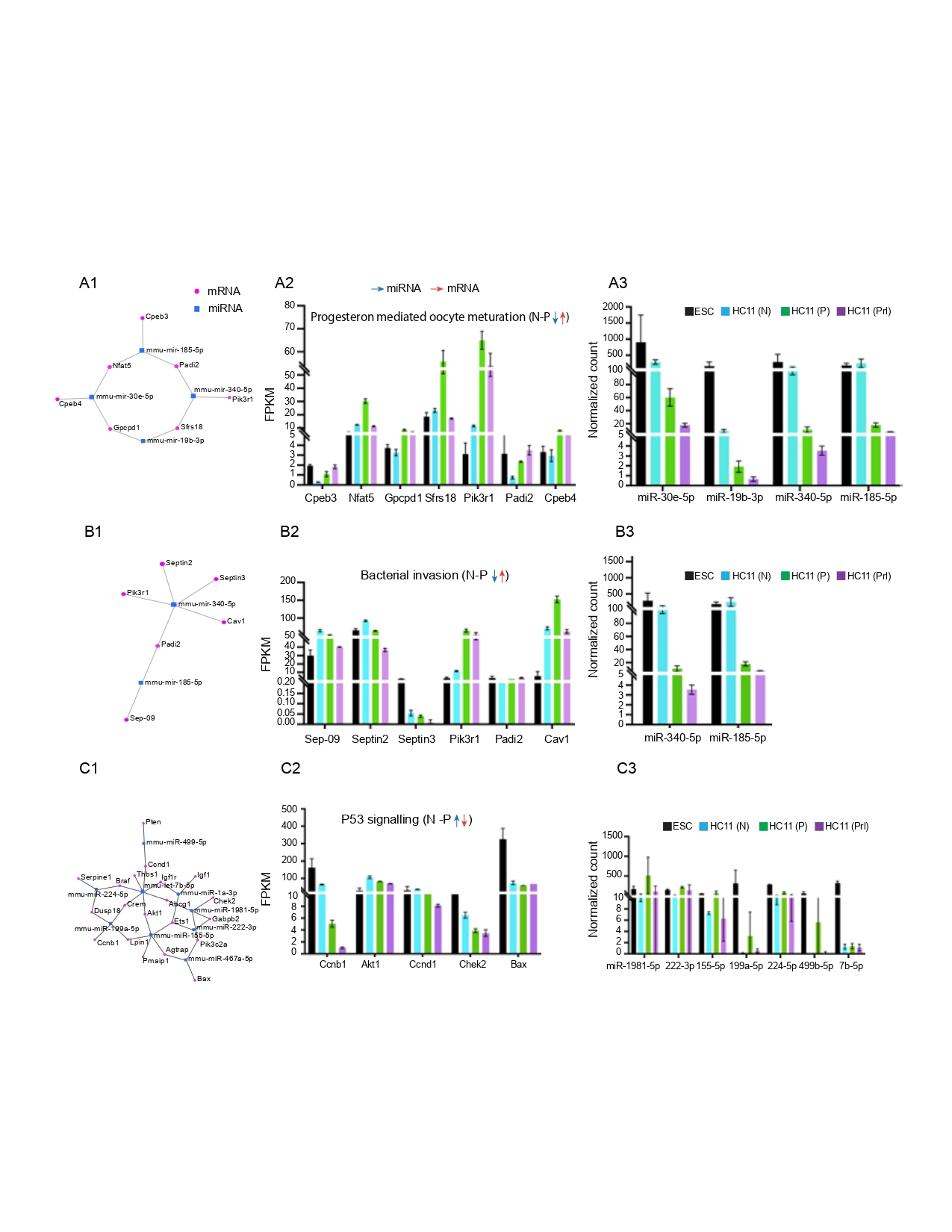

### mRNA-seq and miRNA-seq normalized counts of up- or down-regulated mRNAs and down and up-regulated miRNAs between HC11 (P)-(PRL) cell stages.

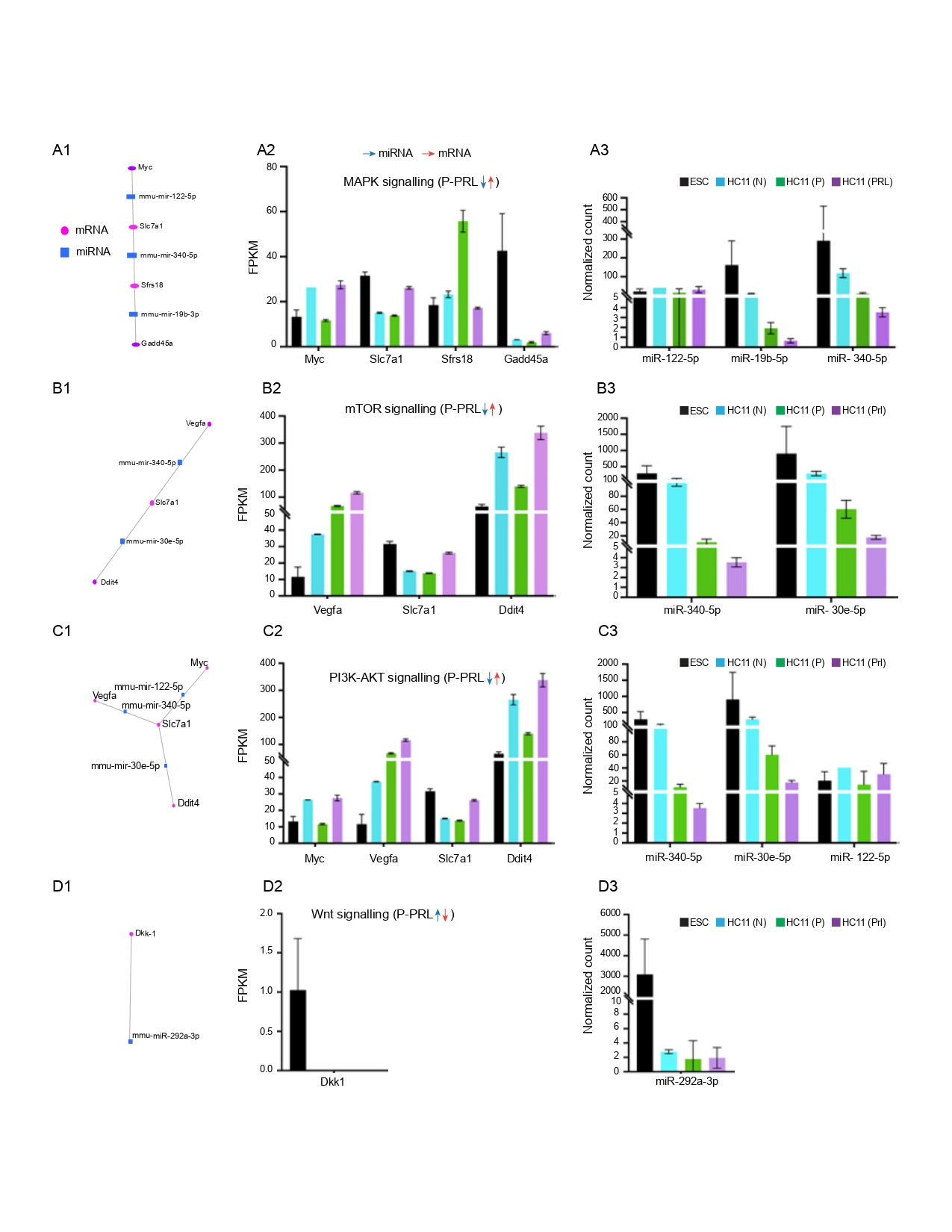

### mRNA-seq and miRNA-seq normalized counts of up-regulated mRNAs and down-regulated miRNAs between HC11 (N)-(P) cell stages.

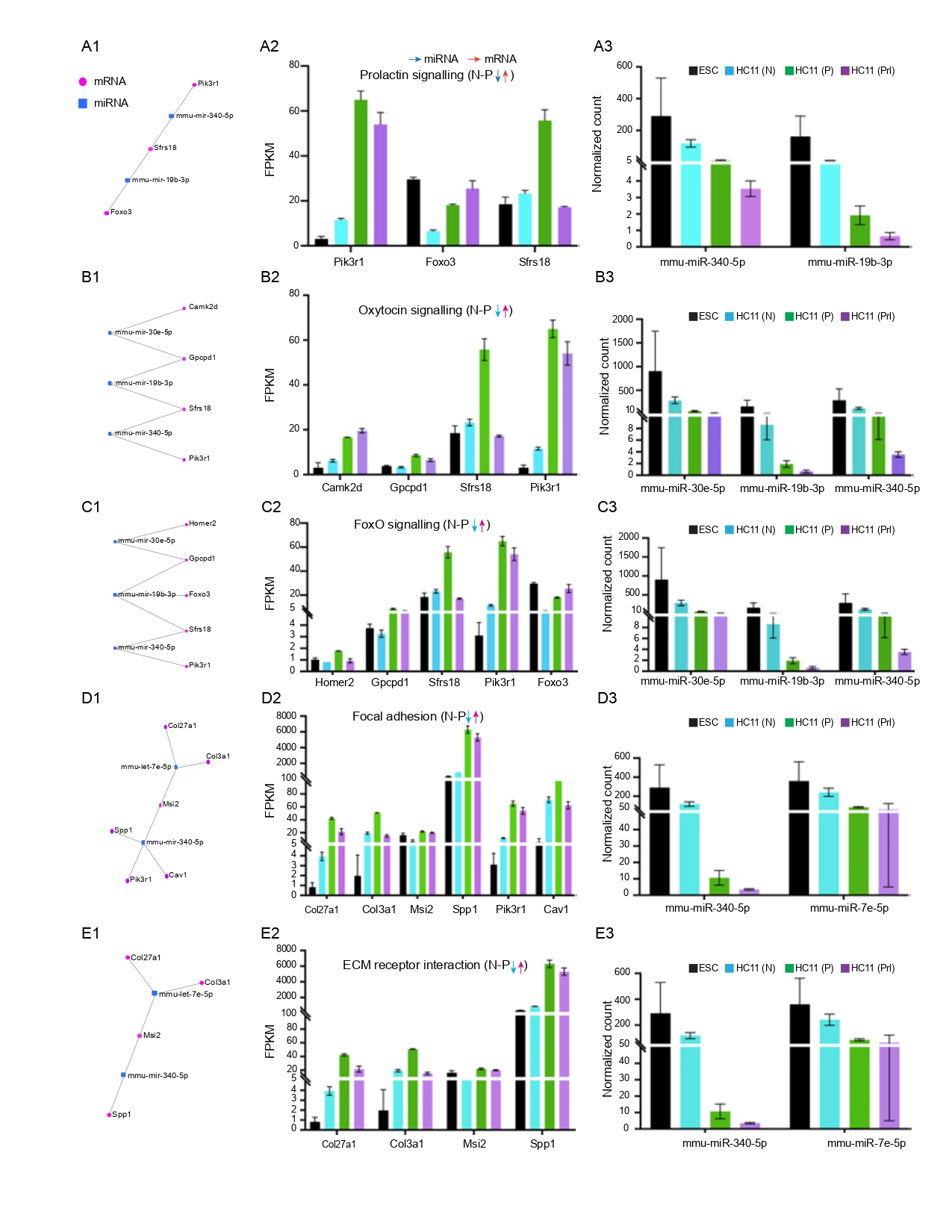

### Statistics of miRNA-sequencing datasets.

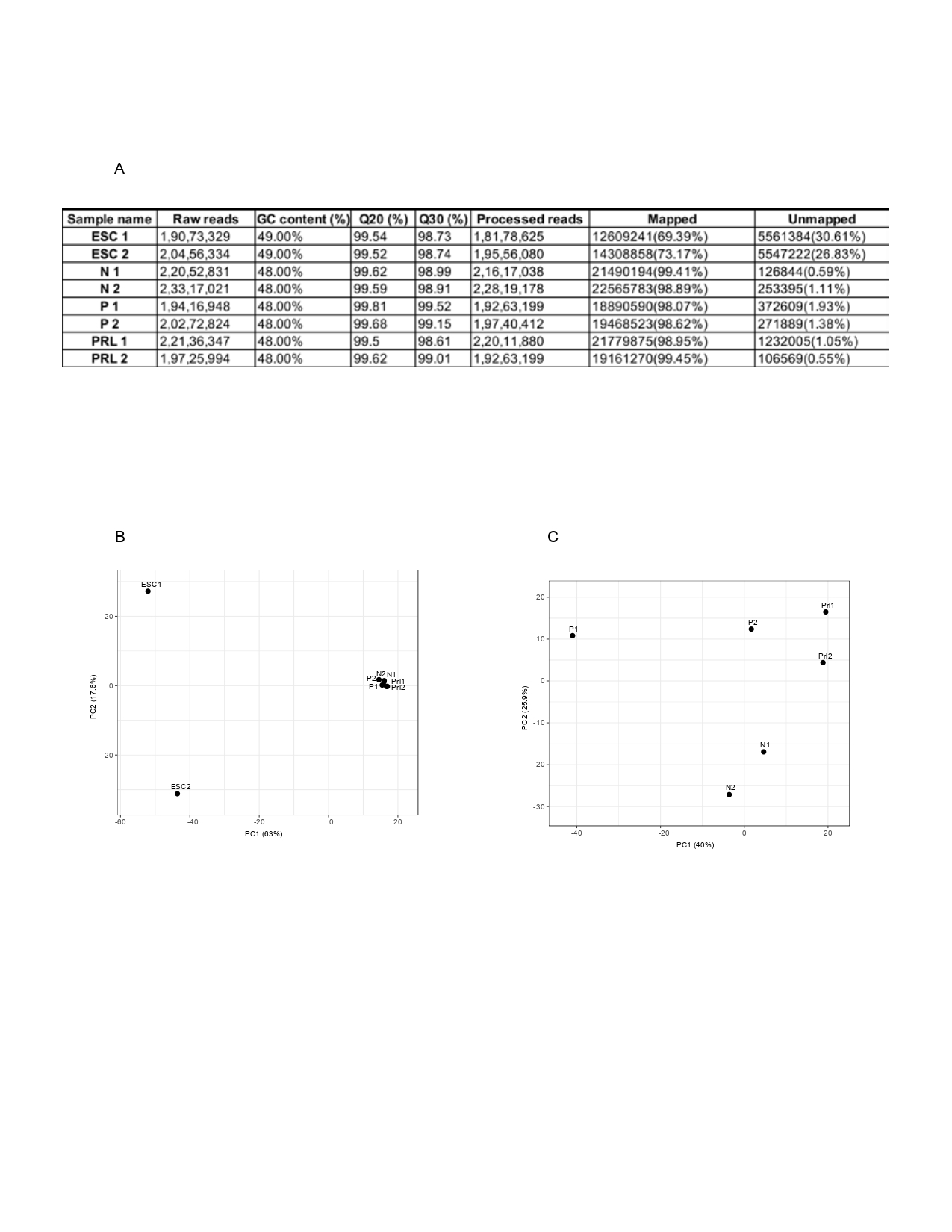
